# Disease swamps molecular signatures of genetic-environmental associations to abiotic factors in Tasmanian devil (*Sarcophilus harrisii*) populations

**DOI:** 10.1101/780122

**Authors:** Alexandra K. Fraik, Mark J. Margres, Brendan Epstein, Soraia Barbosa, Menna Jones, Sarah Hendricks, Barbara Schönfeld, Amanda R. Stahlke, Anne Veillet, Rodrigo Hamede, Hamish McCallum, Elisa Lopez-Contreras, Samantha J. Kallinen, Paul A. Hohenlohe, Joanna L. Kelley, Andrew Storfer

**Affiliations:** School of Biological Sciences, Washington State University Pullman, WA 99164, USA; Plant Biology, University of Minnesota, Minneapolis, MN 55455, USA; Department of Biological Sciences, University of Idaho, Institute for Bioinformatics and Evolutionary Studies, University of Idaho, 875 Perimeter Drive, Moscow, Idaho 83844, USA; School of Biological Sciences, Hobart, TAS 7004, Australia; School of Environment, Griffith University Nathan, QLD 4111, Australia

**Author notes:** Emails for correspondence: Alexandra Fraik, School of Biological Sciences, Washington State University, Pullman, WA,; Andrew Storfer, School of Biological Sciences, Washington State University, Pullman, WA.

**Keywords:** landscape genomics, genetic-environmental associations, Tasmanian devils, devil facial tumor disease, population genetics, local adaptation

## Abstract

Landscape genomics studies focus on identifying candidate genes under selection via spatial variation in abiotic environmental variables, but rarely by biotic factors such as disease. The Tasmanian devil (*Sarcophilus harrisii*) is found only on the environmentally heterogeneous island of Tasmania and is threatened with extinction by a nearly 100% fatal, transmissible cancer, devil facial tumor disease (DFTD). Devils persist in regions of long-term infection despite epidemiological model predictions of species’ extinction, suggesting possible adaptation to DFTD. Here, we test the extent to which spatial variation and genetic diversity are associated with the abiotic environment and/or by DFTD. We employ genetic-environment association (GEAs) analyses using a RAD-capture panel consisting of 6,886 SNPs from 3,286 individuals sampled pre- and post-disease arrival. Pre-disease, we find significant correlations of allele frequencies with environmental variables, including 349 unique loci linked to 64 genes, suggesting local adaptation to abiotic environment. Following DFTD arrival, however, we detected few of the pre-disease candidate loci, but instead frequencies of candidate loci linked to 14 genes correlated with disease prevalence. Loss of apparent signal of abiotic local adaptation following disease arrival suggests swamping by the strong selection imposed by DFTD. Further support for this result comes from the fact that post-disease candidate loci are in linkage disequilibrium with genes putatively involved in immune response, tumor suppression and apoptosis. This suggests the strength GEA associations of loci with the abiotic environment are swamped resulting from the rapid onset of a biotic selective pressure.

## Introduction

A central goal of molecular ecology is understanding how ecological processes generate and maintain the geographic distribution of adaptive genetic variation. Landscape genomics has emerged as a popular framework for identifying candidate loci that underlie local adaptation (Manel et al. 2010; Rellstab et al. 2015; Hoban et al. 2016; Lowry et al 2016; Storfer et al. 2017). Using genomic-scale data sets, researchers screen for loci that exhibit patterns of selection across heterogeneous environments (Haasl et al. 2016). One widely-used method for testing for statistical associations of allele frequencies of marker loci across the genome with environmental variables is genetic-environmental associations (GEAs) (Rellstab et al. 2015; Francois et al. 2016; Whitlock & Lotterhos 2015; Hoban et al. 2016).

GEAs identify significant correlations of allele frequencies at candidate loci with abiotic environmental variables such as altitude, rainfall, and temperature. These efforts have been successful, for example, by identifying hypoxia adaptation in high-elevation human populations (Beall 2007, Peng et al. 2010), stress response in lichen populations along altitudinal gradients (Dal Grande 2017), and leaf longevity and morphogenesis in response to aridity (Steane et al. 2014). Climatic, geographic, and fine-scale remote sensing data that explain large amounts of heterogeneity in the environment are often easily obtained (Rellstab et al. 2015). However, few landscape genomics studies have tested for the influence of biotic variables on the spatial distribution of adaptive genetic variation.

Infectious diseases often impose strong selective pressures on their host and thereby represent key biotic variables increasingly recognized for their severe impacts on natural populations (summarized in Kozakiewicz et al. 2018, e.g. Biek et al. 2010; Wenzel et al 2015; Leo et al. 2016; Mackinnon et al. 2016; Eoche-Bosy et al. 2017). However, data on biotic factors such as life history traits (Sun et al. 2015), community composition (Harrison et al. 2017), or disease prevalence often involve extensive field work and are far more difficult and labor-intensive to collect than abiotic variables. Landscape genomic studies of disease can help guide management programs, such as captive breeding designs and reintroductions (Hoban et al. 2016; Hohenlohe et al. 2018). Additionally, landscape genomics studies have been used elucidate how the landscape influences the distribution of both the host and vector (Blanchong et al. 2008; Robinson et al. 2013; Schwabl et al. 2017). However, studies disentangling the influence of pathogen dynamics from that of other abiotic landscape features have had limited statistical power to date. For example, Wenzel et al. (2015) tested for correlations between parasite burden and genetic differentiation across the landscape but did not detect any statistically significant associations due to noise created by the genomic background. Disease may play an important role in the distribution of adaptive genetic variation across the landscape, potentially swamping signatures of local adaptation to the abiotic environment.

Tasmanian devils (*Sarcophilus harrisii*) and their transmissible cancer, devil facial tumor disease (DFTD), offer such an opportunity. This species is isolated to the island of Tasmania and has been sampled across its entire geographic range. Intense mark-recapture studies and collection of thousands of genetic samples has been conducted over the past 20 years both before and after DFTD arrival across multiple populations. This sampling effort, in addition to resources such as an assembled genome (Murchison et al. 2012), provide extensive data to employ GEAs to test for the relative effects of abiotic environmental factors versus DFTD.

In 1996, the first evidence of DFTD was documented in Mt. William/Wukalina National Park (McCallum 2008). In just over two decades, DFTD has spread across the majority of Tasmania, with nearly a 100% case fatality rate (Hawkins et al. 2006; McCallum et al. 2009). Transmission appears to be largely frequency-dependent (McCallum et al. 2009; McCallum 2012), with tumors originating on the face or in the oral cavity and being transferred as allografts (Pearse et al. 2006) through biting during social interactions (Hamede et al. 2009). DFTD has a single, clonal Schwann cell origin (Siddle et al. 2010) and evades host immune system detection via down-regulation of host major histocompatibility complex (Siddle et al. 2013). Low overall genetic variation in devils resulting from population bottlenecks (Miller et al. 2011) that occurred during the last glacial maximum, as well as during extreme El Niño events 3,000-5,000 years ago (Brüniche-Olsen et al. 2014) has also been attributed to high susceptibility to this emerging infectious disease. DFTD likely imposes extremely strong selection as it threatens Tasmanian devils with extinction with resulting local population declines exceeding 90%, and an overall species-wide decline of at least 80% (Jones et al. 2004; Lachish et al. 2009). However, small numbers of devils persist in areas with long-term infection likely resulting from evolutionary responses in the Tasmanian devil (Epstein et al. 2016; Wright et al. 2017). Indeed, recent work has shown: 1) rapid evolution in genes associated with immune-related functions across multiple populations (Epstein et al. 2016), 2) sex-biased response in a few large-effect loci in survival following infection (Margres et al. 2018a), 3) evidence of effective immune response (Pye et al. 2016) and 4) cases of spontaneous tumor regression (Pye et al. 2016; Wright et al. 2017; Margres et al. 2018b). The latter two studies, however, focused on small geographic areas.

Here, we estimated allele frequencies in 6,886 SNPs both randomly selected and previously shown to be associated with DFTD (Epstein et al. 2016) in 3,286 devils from seven localities throughout Tasmania sampled both prior to and following disease arrival. Our study had five objectives to test the relative effects of abiotic environmental variables and DFTD on spatial genetic variation pre- and post-disease arrival: 1) to estimate the number of genetic clusters to determine whether population declines caused by disease altered population structure; 2) to identify genetic-environmental associations of devil populations to abiotic variables prior to disease arrival; 3) to identify genetic-environmental associations to both abiotic variables and DFTD prevalence post-disease arrival; 4) to determine whether signals of local adaptation to abiotic variables pre-disease were weakened by the strong selection imposed by DFTD; 5) to assess whether changes in genetic variation pre- and post-disease arrival were the result of non-neutral processes, as opposed to genetic drift due to small surviving population sizes. We predict that DFTD epidemics would result in swamping of prior genetic-environmental association with the abiotic environment.

## Methods

### Trapping and sampling data

Field data and ear biopsy samples from 3,286 Tasmanian devils were collected from seven different locations across their geographic range in Tasmania (Figure 1), pre and post-DFTD, between 1999 to 2014 (Table 1). Five of these locations became infected with DFTD during the study; one location remained disease-free, and one location was already infected at the beginning of the study (Figure 1, Table 1). We considered the first year of disease arrival as pre-disease in our analyses, as the generation time for Tasmanian devils is approximately 2 years. Tasmanian devils were trapped using standard protocols (Hamede et al. 2015) involving custom-built polypropylene pipe traps 30 cm in diameter (Hawkins et al. 2006). These traps were set and baited with meat for ten consecutive trapping nights. In each 25 km^2^ trapping site, at least 40 traps were set. Traps were checked daily, commencing at dawn. Each individual was permanently identified upon first capture with a microchip transponder (Allflex NZ Ltd, Palmerstone North, New Zealand). Additional specifics regarding field protocols, samples taken, and phenotypic data recorded can be found in Hamede et al. (2015).

**Figure 1:**
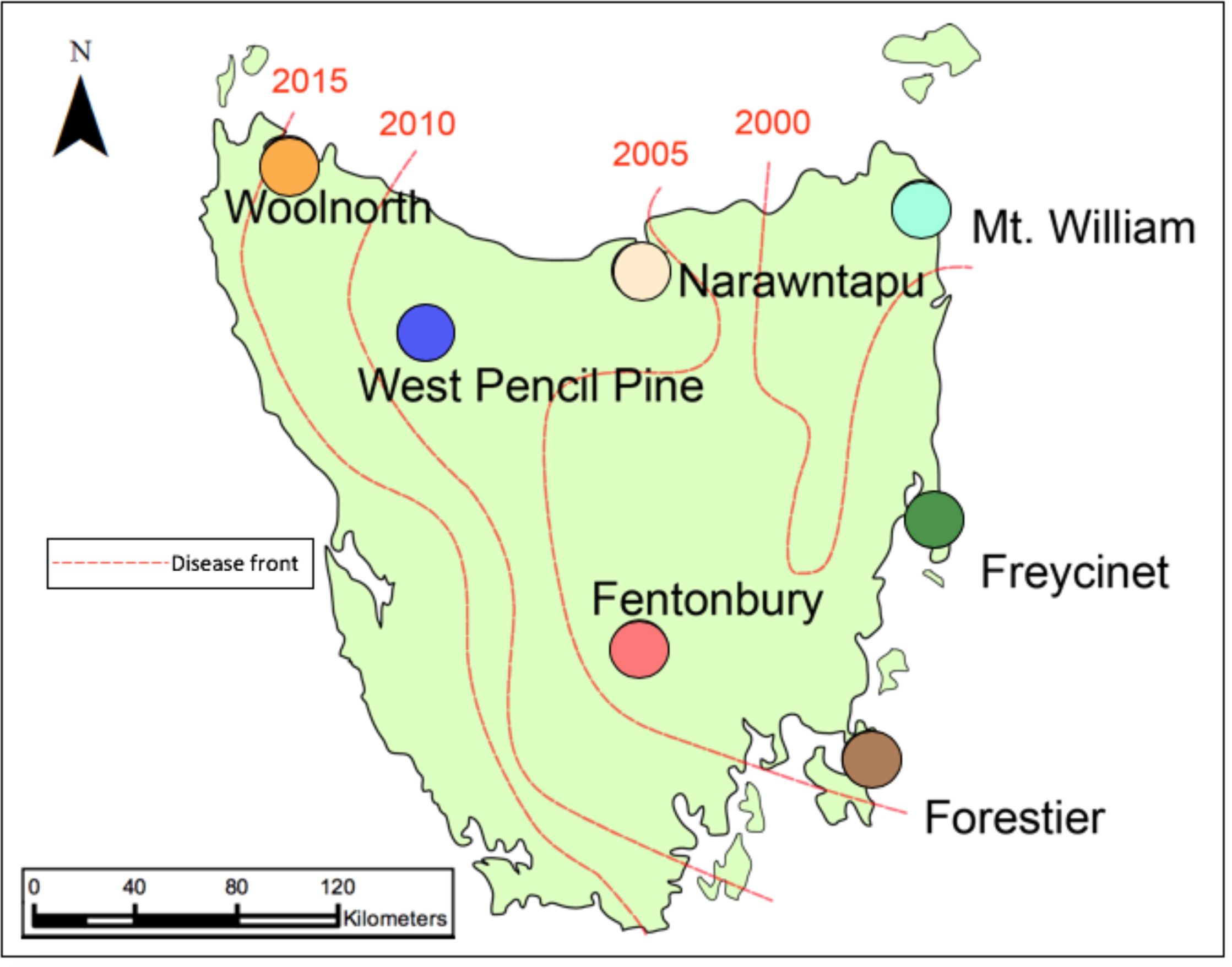
Map of Tasmania with each sampling location. The red lines and the corresponding years indicate the first year that disease was detected in these locations.

**Table 1:** Information for samples collected from each population (the acronyms for the population included in parentheses). Values listed include the number of ear biopsy samples collected per population prior to and post-disease arrival, the years the populations were sampled, and the date that DFTD was first detected in that population.

| **Populations** | **Samples**<br>**Pre-Disease** | **Samples**<br>**Post-Disease** | **Years Samples**<br>**Collected** | **Date of DFTD**<br>**Infection** |
| --- | --- | --- | --- | --- |
| Fentonbury (FEN) | 97 | 157 | 2004-2009 | 2005 |
| Forestier (FOR) | 137 | 382 | 2004-2010,<br>2012-2013 | 2004 |
| Freycinet (FRY) | 511 | 562 | 1999-2014 | 2001 |
| Mt William (MTW) | N/A | 155 | 1999-2014 | 1996 |
| Narawntapu (NAR) | 258 | 153 | 1999-2000, 2003-2012 | 2007 |
| West Pencil Pine (WPP) | 55 | 357 | 2006-2014 | 2006 |
| Woolnorth (WOO) | 491 | N/A | 2006-2010, 20112 | N/A |

### Overview of RAD-capture array development

We used 90,000 Restriction-site Associated DNA sequencing (RAD-seq) loci generated from 430 individuals sampled across 39 sampling localities (Epstein et al. 2016; Hendricks et al. 2017) to develop a RAD-capture probe set (Ali et al. 2015). The details of the sample collection, preparation, and data processing of these original RAD loci is described in Epstein et al. (2016). Using these data, baits were designed to target a total of 15,898 RAD loci. This array included three categories of loci (with some overlap among categories): i) a set of 7,108 loci spread widely across the genome that were genotyped in more than half of the individuals, had ≤ 3 non-singleton SNPs, and had a minor allele frequency (MAF) ≥ 0.05; ii) 6,315 candidate loci based on immune function, restricted to non-singleton SNPs genotyped in ≥ 1/3 of the individuals, in and within 50 kb of an immune related gene; iii) 3,316 loci showing preliminary evidence of association with DFTD susceptibility with ≤ 5 non-singleton SNPs. Each RAD-capture locus was ≥ 20 kb away from other targeted loci to minimize potentially confounding effects of linkage disequilibrium. Additional details of the creation of the RAD-capture probe set have been summarized in Margres et al. (2018a).

### Data quality and filtering

Libraries produced from the RAD-capture arrays were constructed using the *PstI* restriction enzyme and included 3,568 individuals from the seven distinct geographical locations across Tasmania (Figure 1). Libraries were then sequenced on a total of twelve lanes of an Illumina platform (5 lanes on NextSeq at the University of Oregon Genomics & Cell Characterization Core Facility; 7 lanes on HiSeq 4000 at the QB3 Vincent J. Coates Genomics Sequencing Laboratory at the University of California Berkeley). We de-multiplexed paired-end 150 bp reads, removed low quality reads and removed potential PCR/optical duplicates using the clone_filter program (Stacks v1.21; Catchen et al. 2013). We then used Bowtie2 (Langmead & Salzberg 2012) to align reads to the reference genome (Murchison et al. 2012; downloaded from Ensembl May 2017) requiring the entire read align from one end to the other without any trimming (--end-to-end) with sensitive and -X 900 mapping options. With the resulting bam files, we created individual GVCF files using the option to emit all sites in the aligned regions with HaplotypeCaller from GATK (McKenna et al. 2010). All GVCF files were analyzed together using the option GenotypeGVCFs which re-genotyped and re-annotated the merged records. We selected SNPs using the SelectVariants option and filtered using the following parameters: QD < 2.0 (variant quality score normalized by allele depth), FS > 60.0 (estimated strand bias using Fisher’s exact test), MQ < 40.0 (root mean square of the mapping quality of reads across all samples), MQRankSum < -12.5 (rank sum test for mapping qualities of reference versus alternative reads) and ReadPosRankSum < -8 (rank sum test for relative positioning of reference versus alternative alleles within reads). Using VCFtools (Danecek et al. 2011), additional filtering was performed to remove SNPs on the X chromosome, non-biallelic SNPs, indels, those with a minor allele frequency < 0.05, and those missing data at more than 50% of the individuals genotyped and 40% of the sites across all individuals. To parse out the genetic-environmental associations of abiotic factors from disease, samples were divided into pre and post-disease subsets prior to analyses. After filtering, we retained 3,286 Tasmanian devils (1521 before and 1765 after DFTD) and 6,886 SNPs (Table 1).

### Analyses of population structure

To examine the underlying population structure in our dataset, both pre- and post-disease, we used fastSTRUCTURE (Alexander 2009). fastSTRUCTURE is an algorithm that estimates ancestry proportions using a variational Bayesian framework to infer population structure and assign individuals to genetic clusters. We ran both the pre- and post-disease data sets with the number of genetic clusters (K) set from 1 to 18. Genetic relationships amongst sampled populations were also evaluated using discriminant analysis of principal components (DAPC) analysis (Jombart et al. 2010) implemented in the Adegenet package in R (Jombart et al. 2008). This method transforms genotypic data into principal components and then uses discriminant analysis to maximize between group genetic variation and minimize within group variation. To determine the optimal number of genetic clusters, we used the k-means clustering algorithm for increasing values of K from 1 to 20. We selected the optimal K value by selecting the visual “elbow” in the Bayesian information criterion (BIC) score, which is the lowest BIC value that also minimizes the number of components or genetic clusters retained.

### Genetic diversity

To identify signals of a potential genetic bottleneck, we tested for changes in genetic diversity pre- and post-disease emergence. We used VCFtools (Danecek et al. 2011) to calculate Weir and Cockerham’s estimator of F_ST_ (Θ; Weir and Cockerham 1984) for all pairwise comparisons and all pre-post disease population comparisons. To gauge whether populations had significantly different F_ST_ values, we bootstrapped 95% confidence intervals for 10,000 iterations and tested whether the confidence intervals bounded zero. Additionally, we calculated *π* (Nei and Li 1979) and Tajima’s D (Tajima 1989) for each population before and after disease arrival to estimate standing levels of genetic variation (Danecek et al. 2011). Each metric was calculated by taking the average of each non-overlapping windows of 10 kilobases (kb) per population pre and post-disease. Finally, we tested for changes in inbreeding (F_IS_) using the R package Adegenet (Jombart et al. 2008) for each population pre and post-disease.

### Abiotic environmental variables

We used ArcGIS 10 (ESRI 2011) to plot the location of every sampled Tasmanian devil. Over the course of the 15 years (1999-2014; Table 1), there was some variation in the shape of the trapping grids at the sampling sites due to changes in land use and permissions. To account for this variation, we located the centroid of each sampling area using the calculate geometry tool in ArcGIS 10 (ESRI 2011). We then drew a 25 km^2^ ellipse around each centroid to represent the total trapping area for each sampling location and extracted environmental data for each location. Although the Freycinet and Forestier sites were larger than other sampling sites, the centroids were in the middle of the peninsulas likely making them representatives of those sites. The data layers containing these environmental data were obtained from www.ga.gov.au, www.worldclim.org, and www.thelist.tas.gov.au (see Supplementary Table 1 for a list of the 18 abiotic environmental variables analyzed).

We extracted the values for each environmental variable at five randomly chosen locations within the ellipse generated for each trapping area to test whether there was significant heterogeneity for any of the environmental variables across the trapping area of any particular site. Using the calculate statistics tools in ArcGIS 10, we calculated the mean, standard deviation, and 95% confidence intervals for each of the environmental variables within each site. If the centroid fell within the 95% confidence interval of the randomly selected points at each site, we deemed the centroid as an adequate representation for that environmental variable site-wide (Supplementary Table 2).

To estimate collinearity among environmental variables (Supplementary Table 1), we conducted a principal component analysis (PCA) using the prcomp package in R (R Core Team 2013) including the centroid values for each of our seven sampling localities. The top two abiotic environmental variables that explained the greatest proportion of the variance in each of the first six principal components in the PCA were used as explanatory variables in the subsequent GEA analyses. We did not include the seventh principal component as the amount of variation in the data explained by this component was negligible (< 0.001). To ensure that environmental variables were not correlated, we conducted paired correlative tests using the Spearman’s *ρ* statistic.

### Biotic environmental variables

To estimate disease prevalence within populations, we used trapping records compiled in a database provided by collaborators at the University of Tasmania and the Department of Primary Industries, Water and Environment from 1999-2014 from the six infected populations (McCallum et al. 2009; Hamede at al. 2009, 2012, 2013, 2015, unpubl; Lachish et al. 2010). Field records contained data regarding every unique trapping encounter as well as phenotypic data for each devil. As there is currently no early diagnostic test for DFTD, likelihood of a single devil being infected with DFTD is recorded for devils in the field using a scale of 1-5 (Hawkins et al. 2006). We calculated disease prevalence using the total number of unique devils trapped with a visual DFTD score ≥ 3 (indicative of a growth with high probability of being a DFTD tumor) divided by the number of unique devils captured alive that year. In our GEAs, we averaged the annual disease prevalence values across years 2-4 after DFTD was confirmed at a particular sampling location. We selected these years as the trapping and monitoring efforts across these field sites instead of using the first year of infection of DFTD at a particular location as there was variation in estimates of prevalence during the first year of DFTD detection and no guarantee that this truly was the first year of infection.

### Genetic-environmental association (GEA) analyses

We tested correlations of allele frequencies with abiotic and biotic environmental variables across the seven study locations using latent factor mixed models (LEA; Frichot et al. 2015) and Bayenv2 (Gunther and Coop 2013). The R package for landscape and ecological association studies (LEAs) uses latent factors, analogous to principal components in a PCA, to account for background population structure (Frichot et al 2013). These latent factors serve as random effects in a linear model that tests for correlations between environmental variables and genetic variation (Frichot et al 2013). The number of populations sampled was used for the number of latent factors in LEA. Bayenv2 uses allelic data to generate a variance-covariance matrix, or kinship matrix, to estimate the neutral or null model for underlying demographic structure (Coop et al. 2010; Gunther and Coop 2013). Spearman’s *ρ* statistics were then calculated to provide non-parametric rankings of the strength of the association of each genotype with each environmental variable compared to the null distribution described by the kinship matrix alone (Gunther and Coop 2013). The non-parametric Spearman’s rank correlation coefficients have been shown to be more powerful in describing genetic-environmental relationships if there are extreme outliers (Whitlock and Lotterhos 2015; Rellstab et al. 2015).

### Candidate gene identification

Although each of the landscape genomic methods listed above detect GEAs, each program has been shown to vary in true-positive detection rate given sampling scheme, underlying demography, and population structure (Hoban et al. 2016; Lotterhos and Whitlock 2014; 2015; Rellstab et al. 2015). To reduce the false-positive rate without sacrificing statistical power, we used MINOTAUR (Lotterhos et al., 2017; Verity et al. 2017) to identify putative loci under selection. MINOTAUR takes the test statistic output from each of the GEAs and identifies the top outliers in multi-dimensional space; here, we used the Mahalanobis distance metric to ordinate our outliers. We selected this distance metric as it has been shown to have high statistical power on simulated genomic data sets (Verity et al. 2017) and because our data followed a parametric distribution. We chose our final set of candidate SNPs by selecting the top 1% of loci that had the largest Mahalanobis distance for each environmental variable. Thus, we performed seventeen separate MINOTAUR runs: eight for the eight abiotic environmental variables tested pre-DFTD arrival and nine for the eight abiotic and single biotic environmental variable we tested post-DFTD arrival. We performed seventeen MINOTAUR runs to select the top 1% of loci that had the greatest support for associations with each environmental variable across both methods pre and post-disease.

We then used bedtools (Quinlan and Hall 2010) to identify genes within 100 kb of each candidate SNP in the devil reference genome (Murchison et al. 2010). We chose 100 kb as it has been shown to be the conservative estimate of the size of linkage groups in Tasmanian devils (Epstein et al. 2016). Protein-coding genes within these windows were included in the list of candidate genes. Gene annotations were retrieved from the ENSEMBL database (Akey et al. 2016), gene IDs were derived from the NCBI GenBank database (Wheeler et al. 2017), and descriptions of putative function and gene ontologies were gathered from www.genecards.org (Fishilevich et al. 2017) and the Gene Ontology (GO) Consortium (Ashburner et al. 2000). We then tested for enrichment of GO terms in our candidate gene list using the R package SNP2GO (Szkiba et al. 2014). SNP2GO tests for over-representation of GO terms associated with genes within a specified region using Fisher’s exact test and corrects for multiple testing using the Benjamini-Hochberg (1995) false-discovery rate (FDR). We ran this program on the pre- and post-disease candidate sets separately and used all genes within 100 kb of the original RAD-capture data set as our reference set. Enriched terms were those with an FDR ≤ 0.05.

### Fisher’s exact test

To determine if GEAs detected pre-DFTD remained post-DFTD arrival, we calculated the change in the MINOTAUR rank of each candidate locus following disease arrival. We ranked loci by Mahalanobis distance and compared the rank of the pre-DFTD candidate loci to post-disease arrival for each of the eight abiotic environmental variables. To determine whether a greater proportion of loci detected pre-disease had reduced MINOTAUR rankings post-disease arrival than we would expect by random chance, we conducted Fisher’s exact tests using an *α*=0.05. If we detected a significantly higher proportion of models in which the MINOTAUR rankings were lower post-disease than pre-disease, then we considered the molecular signal of the genetic-environmental to be swamped by disease.

## Results

### Population structure and genetic diversity

Our analyses generally supported K=6 pre and post-DFTD arrival reflecting the six populations sampled during these time periods. However, there was uncertainty of the optimal K-value in fastSTRUCTURE. Using the “chooseK” python script recommended in fastSTRUCTURE, K=9 (Supplementary Figure 1a) received the greatest support pre-DFTD and K=8 post-DFTD (Supplementary Figure 1b). However, in both cases, the difference in the marginal likelihood between K=4 and the selected K was very small (pre-DFTD *Δ*Marginal Likelihood=0.003, post-DFTD *Δ*Marginal Likelihood=0.0032; Supplementary Figure 1c,d). For visual comparison, we also plotted K=6 pre-DFTD (Figure 2a) and post-DFTD (Figure 2b), which reflects the number of populations sampled. The K value for DAPC that minimized the BIC, as well as the number of components included, was K=6 both pre-DFTD (Figure 2c) and post-DFTD (Figure 2d). Pairwise F_ST_ values produced using Weir and Cockerham’s estimator would also support K=6 as all bootstrapped confidence intervals for population comparisons did not bound zero (Table 2). We found that there were no significant differences between *p* (Supplementary Table 3), Tajima’s D (Supplementary Table 3), Nei’s F_IS_ (Supplementary Table 3) or Weir and Cockerham’s F_ST_ values for any of the populations after DFTD arrived, suggesting there were no significant changes in genetic diversity following disease arrival.

**Figure 2:**
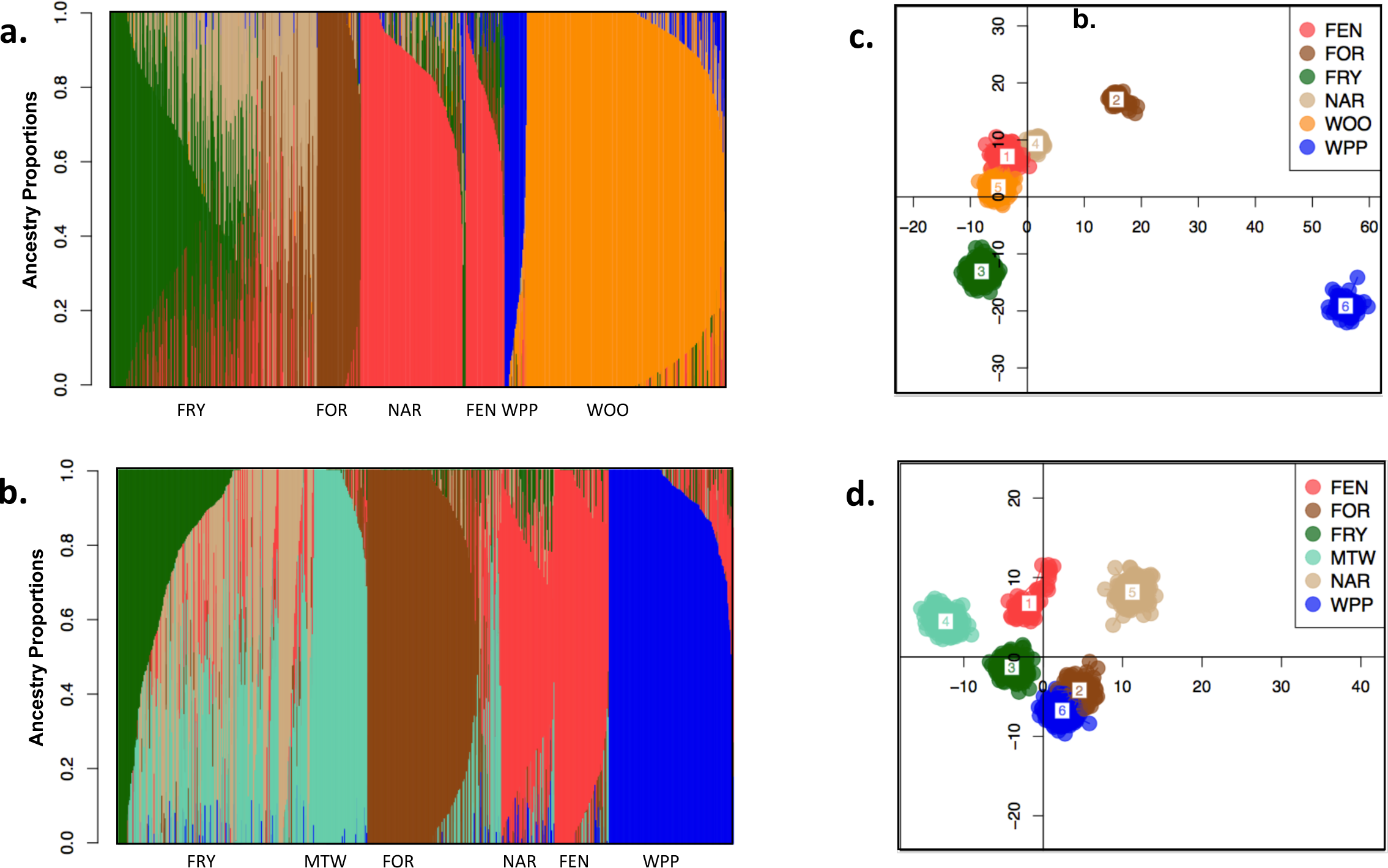
Population assignments computed by fastSTRUCTURE and DAPC for samples collected prior to (a,c) and post (b,d) DFTD arrival. (a,b) Each vertical bar in the fastSTRUCTURE plots represents a single individual sampled at one of the sampling locations which are abbreviated along the x-axis. K=6 is plotted here. Each color in all plots represents a distinct genetic cluster. (c,d) DAPC scatterplots show the first two principal components for K=6.

**Table 2:** The mean pairwise F_ST_ values ± 95% confidence intervals generated by resampling by bootstrapping 10,000 iterations. The values below the diagonal are pairwise comparisons for samples collected pre-disease and those above the diagonal are the same for post-disease. All pairwise comparisons were statistically significant.

|  |  | Post-DFTD |  |  |  |  |  |  |
| --- | --- | --- | --- | --- | --- | --- | --- | --- |
|  |  | FEN | FOR | FRY | MTW | NAR | WOO | WPP |
| Pre-DFTD | FEN |  | 0.049<br>± 0.028 | 0.051<br>± 0.026 | 0.062<br>± 0.035 | 0.026<br>± 0.015 | N/A | 0.066<br>± 0.037 |
|  | FOR | 0.057<br>± 0.033 |  | 0.030<br>± 0.017 | 0.033<br>± 0.019 | 0.0445<br>± 0.025 | N/A | 0.094<br>± 0.052 |
|  | FRY | 0.046<br>± 0.029 | 0.036<br>± 0.021 |  | 0.0212<br>± 0.012 | 0.038<br>± 0.022 | N/A | 0.091<br>± 0.051 |
|  | MTW | N/A | N/A | N/A |  | 0.050<br>± 0.028 | N/A | 0.060 |
|  | NAR | 0.022<br>± 0.013 | 0.050<br>± 0.028 | 0.039<br>± 0.021 | N/A |  | N/A | 0.106<br>± 0.057 |
|  | WOO | 0.156<br>± 0.087 | 0.176<br>± 0.098 | 0.150<br>± 0.080 | N/A | 0.129<br>± 0.072 |  | N/A |
|  | WPP | 0.063<br>± 0.036 | 0.110<br>± 0.059 | 0.104<br>± 0.059 | N/A | 0.061<br>± 0.035 | 0.077<br>± 0.044 |  |

### Environmental variables

We initially considered 18 abiotic environmental variables (Supplementary Table 1) that may be relevant to the distribution of genetic variation across the devil’s geographic range. Eight of these 18 variables were not correlated (Table 3) and explained a significant proportion (> 0.99) of the variance in the environmental data as summarized in the top principal components (Supplementary Table 4). There were no statistically significant differences between the centroid values or any of the five randomly selected points for any of the environmental variables within each of the sampling locations (Supplementary Table 2), indicating a lack of significant within-site heterogeneity. These eight abiotic environmental variables were subsequently used in the GEAs.

**Table 3:**
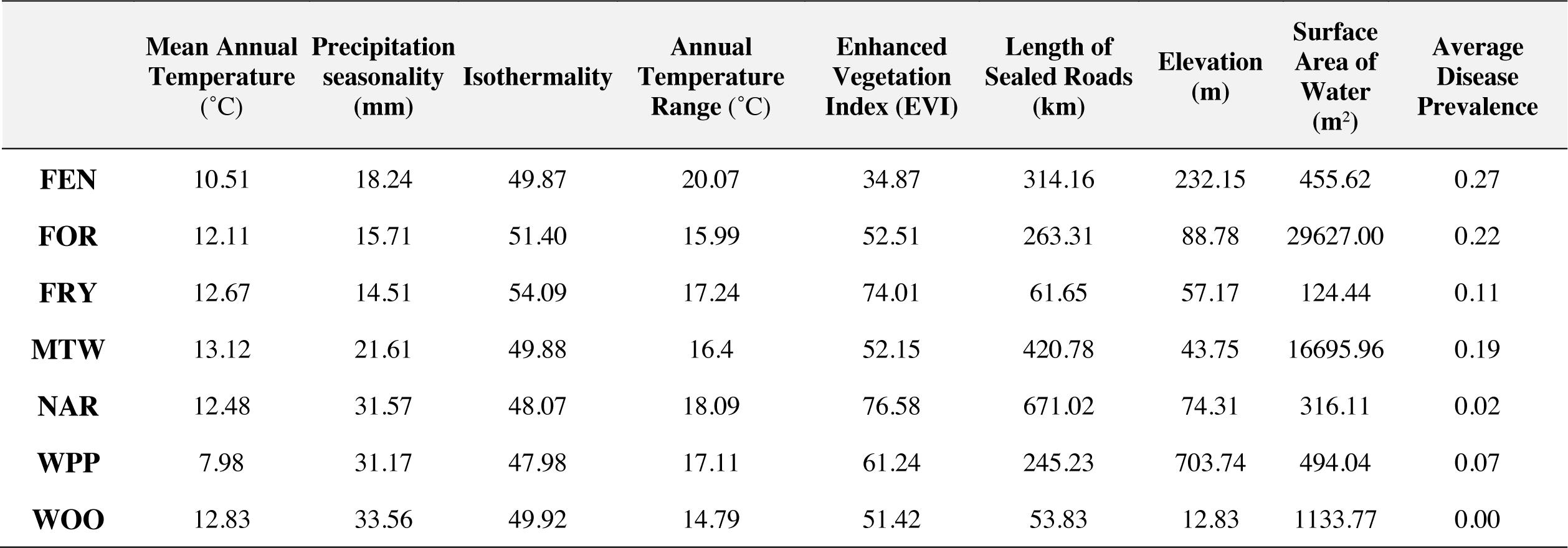
The centroid values for the eight abiotic environmental variables and the biotic variable utilized in the GEA associations for each population.

|  | Mean Annual<br>Temperature<br>(°C) | Precipitation<br>seasonality<br>(mm) | Isothermality | Annual<br>Temperature<br>Range (°C) | Enhanced<br>Vegetation<br>Index (EVI) | Length of<br>Sealed Roads<br>(km) | Elevation<br>(m) | Surface<br>Area of<br>Water<br>(m<sup>2</sup>) | Average<br>Disease<br>Prevalence |
| --- | --- | --- | --- | --- | --- | --- | --- | --- | --- |
| **FEN** | 10.51 | 18.24 | 49.87 | 20.07 | 34.87 | 314.16 | 232.15 | 455.62 | 0.27 |
| **FOR** | 12.11 | 15.71 | 51.40 | 15.99 | 52.51 | 263.31 | 88.78 | 29627.00 | 0.22 |
| **FRY** | 12.67 | 14.51 | 54.09 | 17.24 | 74.01 | 61.65 | 57.17 | 124.44 | 0.11 |
| **MTW** | 13.12 | 21.61 | 49.88 | 16.4 | 52.15 | 420.78 | 43.75 | 16695.96 | 0.19 |
| **NAR** | 12.48 | 31.57 | 48.07 | 18.09 | 76.58 | 671.02 | 74.31 | 316.11 | 0.02 |
| **WPP** | 7.98 | 31.17 | 47.98 | 17.11 | 61.24 | 245.23 | 703.74 | 494.04 | 0.07 |
| **WOO** | 12.83 | 33.56 | 49.92 | 14.79 | 51.42 | 53.83 | 12.83 | 1133.77 | 0.00 |

### Landscape genomics analyses

Combining GEAs across all abiotic variables, we identified 349 unique loci pre-DFTD and 397 unique loci post-DFTD that received the greatest support from MINOTAUR as candidate loci.

We identified candidate genes based on linkage disequilibrium (within 100 kb) to the unique loci, which resulted in 64 genes pre-disease (Supplementary Table 5) and 83 genes post-disease (Supplementary Table 6); seven of those genes overlapped temporally (Supplementary Figure 2, Supplementary Table 5). There were 58 additional unique loci detected by MINOTAUR post-DFTD arrival as related to disease prevalence in the GEAs. Within 100 kb of these 58 SNPs, there were seven annotated genes, none of which overlapped temporally with the pre-disease candidate set (Supplementary Table 6). For candidate genes in which there was more than a single association with a particular environmental variable, the variable that had the greatest absolute Pearson’s correlate value was selected to be used in these candidate gene-environment associations.

### Candidate gene identification for selection by the abiotic environment

Mean annual temperature and annual temperature range were the two most common variables significantly correlated with the allele frequencies of candidate SNPs amongst devil populations pre-disease arrival (Supplementary Table 7). Twenty-nine of the top 64 candidate genes pre-disease were associated with at least one of these two environmental variables and broadly associated with response to external stress and ion transport (Supplementary Tables 5, 7).

Specifically, genes associated with annual temperature range had GOs including RNA transport and processing, and gene pathway regulation. Mean annual temperature, in contrast, was correlated with genes with intracellular protein transport, protein and lipid binding, and cell adhesion GOs. Additionally, isothermality was frequently correlated with significant differences in allele frequency of candidate loci among populations and have similar GOs as annual temperature range. Elevation and vegetation index explained a large proportion of the observed variance in the abiotic environment across our seven sampling sites (Supplementary Table 2). Elevation was correlated with seven of the candidate genes with ion binding and oxidoreductase activity functions (Supplementary Tables 5, 7). Vegetation index was correlated with allele frequencies of thirteen of the candidate genes that had GOs including transcription and cellular protein binding.

In our abiotic GEAs post-disease arrival annual temperature range, mean annual temperature and vegetation index were the top environmental variables most frequently associated with candidate SNPs (Supplementary Tables 6, 8). Similar to pre-DFTD, GOs for genes associated with these two temperature variables included cellular response to stimulus, RNA processing and regulation, and ion binding. In contrast to pre-DFTD, mean annual temperature and annual temperature range were associated with genes with cell signaling and apoptotic processes. Vegetation index was associated with fourteen of the candidate genes post-DFTD with GOs including transcription and cell signaling pathways (Supplementary Tables 6, 8).

There was no statistically significant enrichment of any GO category of our candidates (pre-disease p=0.344; post-disease p=0.297), there were commonalities in the putative functions of candidate genes pre- versus post-disease including regulation of transcription and translation and cellular response to external stress (Supplementary Tables 5, 6).

### Candidate gene identification for selection by the biotic environment

Disease prevalence was significantly correlated with divergent allele frequencies among sampled devil populations (Supplementary Table 8, 9). Fourteen of the 64 candidate genes detected post-DFTD arrival were associated with disease-prevalence (Supplementary Tables 8, 9). The associated GOs for these fourteen genes included tumor regression, apoptosis, and immune response (Supplementary Table 6). Of these fourteen genes, six genes were also associated with abiotic environmental variables. Genes *TBXAS1* and *FGGY* were found in both the pre and post-DFTD candidate gene lists and associated with GOs including oxidoreductase activity and metabolism (Supplementary Table 5).

### Fisher’s exact test for disease swamping

Putative GEAs found pre-DFTD were consistently undetected post-DFTD arrival (Supplementary Tables 5,6). Using Fisher’s exact tests, fewer pre-DFTD candidate loci were detected post-DFTD arrival than expected by random chance for each MINOTAUR analysis for each of the eight abiotic environmental variables (Figure 3, Supplementary Figure 3). For example, we only detected three of the putative pre-DFTD candidate loci using MINOTAUR post-DFTD arrival for genetic associations with mean annual temperature across sampled populations (Fisher’s Exact Test Odd’s ratio=23.211, p < 0.001; Figure 3).

**Figure 3:**
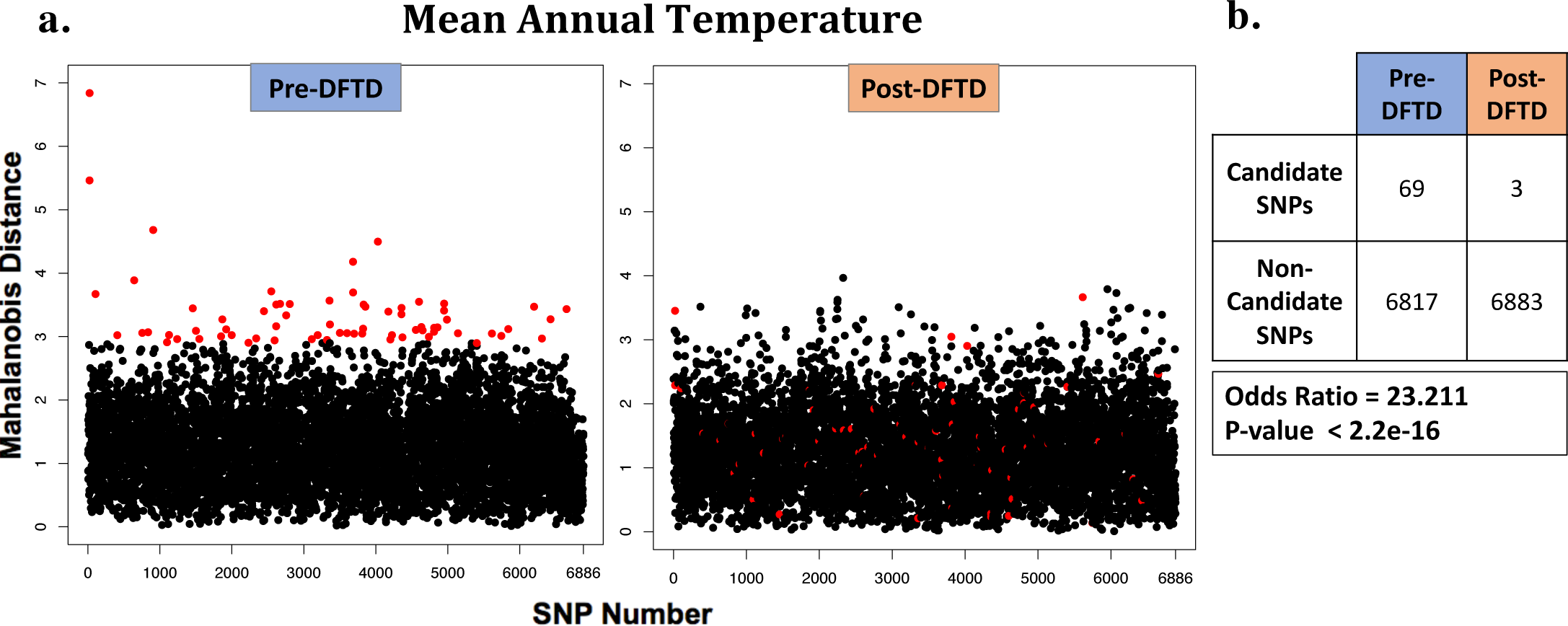
Mahalanobis distances from MINOTAUR for each SNP pre-DFTD arrival (left plot) and post-DFTD arrival (right plot) for GEAs with mean annual temperature. SNPs are ordered by position along the chromosomes. The top 1% of loci with the largest Mahalanobis distance values pre-DFTD are indicated in red. (b) Post-hoc analyses found that a significantly greater number of candidate loci detected pre-DFTD arrival were not detected post-DFTD arrival. This trend was detected consistently across all eight of the abiotic environmental variables tested in genetic-environmental association analyses (output from the remaining seven variables found in supplementary figure 2).

Seven of the original 64 candidate genes detected pre-disease overlapped with the 83 candidate genes post-disease arrival (Supplementary Table 5). Four were associated with at least one of the same environmental variables post-disease arrival as pre-disease, and three were correlated with different variables (Supplementary Table 9). Two of the variants linked with *TBXAS1* and *FGGY* were found to be associated with disease prevalence post-disease arrival. For example, the *FGGY* gene was originally associated with mean annual temperature prior to disease arrival, but disease prevalence post-arrival. Putative functions of FGGY include carbohydrate phosphorylation and neural cell homeostasis (Singh et al. 2017; Dunckley et al. 2007; Supplementary Table 5). Similarly, *TBXAS1*, which is putatively involved in oxidoreductase activity (Ullrich et al. 1994), was associated with mean annual temperature pre-disease arrival, but was more strongly correlated with disease prevalence following disease arrival (Supplementary Tables 5, 9).

## Discussion

Here, we showed Tasmanian devils are more genetically structured across Tasmania than previously documented (Bruniche-Olsen et al. 2014; Hendricks et al. 2017; Miller et al. 2011) thereby creating conditions that favor local adaptation. Indeed, GEAs showed signatures of selection across populations, suggestive of adaptation to local abiotic factors prior to disease arrival. Mean annual temperature and annual temperature range were abiotic variables most frequently associated with candidate genes (Loyau et al. 2016; Tseng et al. 2011; Xu et al. 2017), and these abiotic factors have been established as important for devil habitat use (Jones and Barmuta 2000; Jones and Rose 1996). Isothermality and vegetation index were also associated with candidate genes and correlated with the largest amount of geographic heterogeneity across the landscape (Schweizer et al. 2016; Mukherjee et al. 2018). Nonetheless, most of the candidate genes detected prior to disease arrival were not detected post-DFTD emergence reflecting the strong selection imposed by DFTD. Instead, GEAs showed evidence of significant correlations of disease prevalence with allele frequencies at several SNPs after disease arrival. Taken together, these results suggest that DFTD swamps molecular signatures of local adaptation to abiotic variables. Finally, despite large declines in census population size across all diseased populations, no significant changes in genetic diversity were detected, suggesting detected patterns were not likely the result of genetic drift.

### Genetic population structure

We found conflicting results among our demographic analyses of our sampled populations. Similar to previous studies, we detected admixture between some of the seven sampling locations using fastSTRUCTURE (Figure 2a), which showed an optimal K value between 4-9. However, using DAPC (Figure 2b), we detected distinct genetic clusters among the sampling locations, irrespective of disease presence. Pairwise population F_ST_ calculations were all significantly greater than 0, indicating significant population differentiation between all populations sampled (Table 2). Regardless of the analysis, we identified greater amounts of genetic structure than previously detected. Two previous studies found high levels of admixture between two genetic clusters separating eastern and western Tasmania (Brüniche-Olsen 2014; Hendricks et al. 2017), while another, based on mtDNA genomes, suggested three genetic clusters (Northwest, East and Central; Miller et al. 2011). Our results differed from the previous work because we included much larger sample sizes and numbers of loci, which likely increased our power to detect population structure.

### Genetic diversity

Despite extensive, range-wide population declines, we found no evidence for loss of genetic diversity following disease arrival. Devil populations had low levels of *π* across the species’ geographic range, regardless of whether DFTD was present (Supplementary Table 6), consistent with findings from previous studies (Bruinche-Olsen et al. 2014; Hendricks et al. 2017; Jones et al. 2004). While there was population genetic differentiation (F_ST_) between sampled populations, there was no significant difference in inbreeding coefficient (F_IS_) among populations pre-versus post-disease arrival. Additionally, there was no significant change in genetic differentiation (F_ST_) temporally among populations post-DFTD arrival. Lack of a significant change in the positive Tajima’s D values across populations following disease arrival (Supplementary Table 3) also suggests possible maintenance of some levels of standing genetic variation even after substantial population declines. One possibility for the lack of detectable changes in genetic diversity would be that simply not enough time has passed since the bottleneck. If we take account of the increased precocial breeding in diseased populations, which reduces the generation time of females from 2-3 years prior to disease outbreak to 1.5-2 years (Jones et al. 2008) the maximum number of generations in our data set since disease arrival would be in Freycinet for 8-9 generations (number of generations passed for all populations in Table 1). Detection of significant changes in genetic diversity requires substantial reduction in effective population size for several generations (Luikart et al. 2010). Following DFTD outbreak, devil populations typically decline by 90% after 5-6 years. This time lag, coupled with the short timescale, perhaps resulted in low power to detect changes in genetic diversity.

### Landscape genomics pre-disease

The abiotic environmental variables used in our GEAs have been shown to be important determinants of devil distribution (Jones and Barmuta 2000; Jones and Rose 1996; Rounsevell et al. 1991). Devils are distributed throughout the diverse habitats of Tasmania, but their core distribution comprises areas of low to medium rainfall which are dominated by dry, open forest, woodland and coastal shrubland. They use varying vegetation types across the environment for different functions. Devils prefer a clear understory for movement, forest-grassland edges for hunting, and generally avoid structurally complex vegetation and landscape features such as rocky areas and steep slopes (Jones and Barmuta 2000; Jones and Rose 1996). Candidate loci linked to the CUX1 gene were strongly associated with vegetation index in devils, especially in heavily forested populations such as WPP (Supplementary Table 7). CUX1, which is putatively involved in transcription regulation (Sansregret 2008) and limb development in morphogenesis (Lizarraga et al. 2002), has been detected as a candidate in the mammalian systems for adaptation to inhabiting environments with complex understory (Schweizer et al. 2016; Mukherjee et al. 2018). Landscape genomic analyses of grey wolves found evidence for the involvement of this gene, and others related to auditory function and morphogenesis, in navigating between dense and more open landscape habitat types (Schweizer et al. 2016). Differential expression of this gene in muscle tissue of the bovine species *Bos frontalis* relative to domestic cattle was suggested to assist in reducing energetics required for navigating complex, hilly environments in India (Mukherjee et al. 2018). Strong genetic-environmental association across devil populations throughout their heterogeneous geographic range provide evidence for local adaptive genetic variation.

Although we found no significant enrichment for any GO category in the 64 genes identified in strong LD with the top 1% of SNPs pre-DFTD, 29 of the genes were putatively involved in stress response and ion binding. Climatic variables including mean annual temperature, annual temperature range, and isothermality were all important in describing observed genetic variation across the devil geographic range. The *RPL7L1* gene, for example, which functions in mRNA transcription and translation during mammalian cellular development and differentiation (Maserati et al. 2012), was strongly associated with annual temperature range. In Chinese Sanhe cattle, *RPL7L1* was found to be down-regulated when exposed to severe cold, reflective of an adaptive response to rapid temperature variation (Xu et al. 2017). In devils the allele frequencies of candidate loci linked to this gene appear to be strongly associated with populations occupying the narrowest temperature ranges in devils’ geographic range including NAR, WPP and FRY (Supplementary Table 1). Expressional changes of this gene may therefore be a crucial local adaptation to devils living in these habitats when temperature fluctuates outside the normal range. Additionally, mean annual temperature was significantly associated with genes such as *CYP39A1*, which putatively functions in oxidoreductase activity of steroids and lipid metabolism (Li-Hawkins et al. 2000). This gene is differentially expressed in response to long-term temperature variation, for example, it is downregulated in chicken embryos following extended heat exposure (Loyau et al. 2016) and upregulated in zebrafish brains in response to severe cold (Tseng et al. 2011). Overall, the functions of genes associated with the abiotic variables would suggest specific, local responses to some key environmental conditions that explain devil geographic distributions.

### Landscape genomics pre- vs post-disease

Following the arrival of DFTD, both the number of loci and proportion of variation in the allele frequencies of the pre-disease candidate loci explained by the abiotic environment were significantly reduced. The *TMCC2* gene, for example, was strongly correlated with mean annual temperature prior to disease arrival (pre-disease, Pearson’s correlate=0.915, MINOTAUR rank=39), but the strength of the genetic-environmental association significantly decreased post-disease arrival and it was no longer detected as a candidate locus (post-disease, Pearson’s correlate=-0.3, MINOTAUR rank=6071). Only seven of the 64 candidate genes detected pre-disease were included in the 83 genes detected post-disease set, with only one gene, *ARPC2* (Actin related protein 2/3 Complex Subunit 2), correlated with the same variable prior to disease arrival. *FGGY*, for example, was strongly correlated with mean annual temperature prior to disease arrival (pre-disease, Pearson’s correlate=-0.745, MINOTAUR rank=018). Post-disease arrival, the strength of this relationship significantly decreased (post-disease, Pearson’s correlate=-0.082, MINOTAUR rank=6199), and the SNP linked to this gene was more strongly correlated with disease prevalence (post-disease, Pearson’s correlate=-0.082, MINOTAUR rank=6199). These discordant patterns of association of candidate loci with environmental variables pre compared to post-disease arrival may possibly be explained by pleiotropy or loss of association due to DFTD.

We detected 83 candidate genes post-disease arrival associated with both abiotic environmental variables and/or disease prevalence. Although there was no significant enrichment of any GO category, most genes associated with abiotic variables had stress response GOs, whereas post-disease many genes had GOs involved in cellular processes including apoptosis, cell differentiation and cell development, similar to what has been previously detected (Epstein et al. 2016; Margres et al. 2018a; Frampton et al. 2018). Genes involved in apoptosis detected in our GEAs, including *DNAJA3* in the heat shock protein family, have also been previously identified in DFTD literature as candidates for anti-cancer vaccines (Tovar et al. 2018) for their putative involvement in immunogenicity (Graner et al. 2000) and tumor suppression (Shinagawa et al. 2008). Functionally similar to *PAX3* a gene associated with apoptosis and angiogenesis (Asher et al. 1996) and devil tumor regression detected in previous studies (Wright et al. 2017), we detected genes including *FLCN* and *BMPER* post-DFTD arrival (Qi et al. 2009). We also identified *MGLL* and *TLR6* in our post-DFTD candidate list which are involved in inflammatory response (Epstein et al 2016; Margres et al. 2018a).

The identification of candidate genes putatively involved in immune response (e.g. *RFN126* and *IL9R*), tumor suppression (e.g. *DMBT1*, *FLCN* and *ITFG1*), signal transduction (e.g. *ARHGEF37* and *PPP1R12B*) and cell-cycle regulation (e.g. *TK2*) associated with disease prevalence post-DFTD arrival provide evidence that disease may have had a strong influence on standing genetic variation among devil populations (Tsai et al. 2014; Purwar et al. 2012; Mollenhauer et al. 1997; Hasumi et al. 2015; Leushacke et al. 2011; Tsapogas et al. 2003; Bannert et al. 2003; Sun, Eriksson & Wang et al. 2014).

The lack of GO enrichment may be an artifact of the RAD-capture panel. Loci targeted herein were linked to or found in coding regions of both putatively neutral genes as well as those involved in immune response, potentially biasing our GO enrichment analysis. Another explanation for the lack of enrichment could stem from our conservative approach for identifying outliers. Using MINOTAUR, we took the top 1% of loci from each GEA as our putative candidates. Although this approach reduces false positives, it also can reduce true positives and imposes an upper bound on the number of possible associations we can ascertain between our candidate loci and disease.

### Support for disease swamping abiotic genetic-environmental associations

Intense, long-term monitoring of Tasmanian devils coupled with an expansive temporal and geographic dataset provided the unique opportunity to test for changes in statistical signatures of GEAs. We hypothesized that DFTD would serve as an extreme selective event as it is nearly 100% lethal (Hamede et al. 2015) and produces a rapid adaptive response (Epstein et al 2016; Wright et al. 2017; Margres et al. 2018a,b) that could swamp signals of genetic-environmental associations with the abiotic environment found pre-disease. Our comparisons of candidate loci detected pre- and post-disease support this hypothesis. Although annual temperature range and vegetation index continued to be important drivers of selection in devil populations regardless of disease presence, most candidate genes identified pre-DFTD were not correlated with these variables after DFTD arrived. (Figure 3; Supplementary Figure 3).

The observed loss of pre-DFTD GEAs following disease arrival could also be the result of stochastic processes operating on our populations following a significant demographic event (Lande 1976; Lande 1993; Shaffer 1981). Following large catastrophes, locally-adapted genotypes may be displaced or swamped out randomly by new genetic variation from neighboring populations via genetic rescue (Waddington 1974; Brown and Kodric-Brown 1977). If this were the case, our observation of loss of pre-DFTD candidate genes may be due to drift resulting from demographic change induced by DFTD versus the selection that DFTD imposed.

However, numerous studies showing evidence of the rapid evolutionary response to DFTD across devil populations suggest that DFTD is selecting for particular genetic variants post-DFTD arrival (summarized in Russell et al. 2018 & Storfer et al. 2018). Additionally, in spite of large decreases in population size, there have been no detectable significant changes to genetic diversity following disease arrival.

Traditional landscape genomics studies incorporating biotic variables have successfully identified signatures of adaptation to biotic variables, such as predators (Ritcher-Boix et al. 2011), parasites (Orsini et al. 2013; Wenzel et al. 2016) and symbionts (Harrison et al. 2017). However, few have successfully identified GEAs with infectious disease using the landscape genomics framework (reviewed in Kozakiewicz et al. 2018). In this study we identified genes putatively associated with an emerging infectious disease and quantified how the onset of disease reduced evidence of GEAs with the abiotic environment among populations of Tasmanian devils across their geographic range. Here, we document a novel, biotic variable swamping out molecular signals of association to the abiotic environment. The findings of this study demonstrate the utility of landscape genomics as a tool to explicitly test the influence of biotic factors, such as disease, on the spatial distribution of genetic variation.

## Conclusion

Emerging infectious diseases are increasingly recognized as significant threats to biodiversity and in extreme cases, species’ range contractions and extinctions (Smith et al. 2006). Yet, few landscape genomics studies have tested for statistical correlations of allele frequencies with biotic variables such as infectious diseases. Here, we show that a lethal disease of Tasmanian devils apparently swamps associations of abiotic variables with allele frequencies in devil populations detected before disease arrival. This finding is consistent with previous work that showed rapid evolution of Tasmanian devils in response to disease (Epstein et al. 2016; Wright et al. 2017; Margres et al. 2018a,b). Additionally, no appreciable declines in genetic diversity were detected across multiple analysis methods. Taken together, these results suggest that observed patterns of allele frequency correlations with disease prevalence are more likely attributable to selection than genetic drift. Our approach and results suggest that future landscape genomics studies may prove valuable for identifying loci under selection pre- and post-disease.

## Acknowledgements

We thank Omar Cornejo, and his lab, David Crowder, and Joanna Kelley’s lab for their helpful review and insightful discussion. We are grateful to the anonymous reviewers for feedback on our manuscript. This work was funded by NSF grant DEB-1316549 and NIH grant R01-GM126563 as part of the joint NSF-NIH-USDA Ecology and Evolution of Infectious Diseases program (AS, PAH, MJ and HM).

## Author Contributions

AKF conducted the analyses and wrote the paper; MJM, SB and BE assisted with the analyses and helped write the paper; BS, SH, AV, and AS assisted with the sample preparation and helped write the paper; RH, and MJ collected the samples, contributed to the study design, managed field surveys and helped write the paper; HM helped write the paper; EL-C, and SJK assisted with the analyses; PH, JLK, and AS directed the project, supervised this work and helped write the paper.

## Ethics

Animal use was approved under IACUC protocol ASAF#04392 at Washington State University.

## Data Accessibility

The sequence data has been deposited at NCBI under BioProject PRJNA306495 (http://www.ncbi.nlm.nih.gov/bioproject/?term=PRJNA306495).

Code is available at github.com/jokelley/devil-landscape-genomics.

