## Supplementary_Tables for "Disease swamps molecular signatures of genetic-environmental associations to abiotic factors in Tasmanian devil (*Sarcophilus harrisii*) populations"

**Supplementary Table 1**: The eighteen abiotic environmental variables used in the landscape genomic analyses. Values are taken from the estimated centroid for each sampling location.

| | Fentonbury (FEN) | Forestier (FOR) | Freycinet (FRY) | Mt. William (MTW) | Narawntapu (NAR) | West Pencil Pine (WPP) | Woolnorth (WOO) |
|---|---|---|---|---|---|---|---|
| Annual Rainfall (mm) | 699.38 | 777.36 | 702.18 | 822.80 | 814.25 | 221509.00 | 1069.33 |
| Annual Temperature Range (˚C) | 200.75 | 159.86 | 172.44 | 163.89 | 180.86 | 171.09 | 147.90 |
| Coded Total Tree Cover | 31.22 | 55.12 | 73.70 | 49.12 | 76.59 | 61.84 | 49.40 |
| Elevation ($m^2$) | 232.15 | 88.78 | 57.17 | 43.75 | 74.31 | 703.74 | 12.83 |
| Enhanced Vegetation Index (EVI) | 34.87 | 52.51 | 74.01 | 52.15 | 76.58 | 61.24 | 51.42 |
| Isothermality | 49.87 | 51.40 | 54.09 | 49.88 | 48.07 | 47.98 | 49.92 |
| Maximum Annual Temperature (˚C) | 16.75 | 16.45 | 17.55 | 16.80 | 17.00 | 11.90 | 17.00 |
| Mean Annual Temperature (˚C) | 105.09 | 121.05 | 126.72 | 131.18 | 124.84 | 79.81 | 128.33 |
| Mean Diurnal Temperature Range (˚C) | 100.63 | 83.01 | 94.25 | 82.21 | 87.90 | 83.03 | 74.20 |
| Mean Temperature Coldest Quarter (˚C) | 60.13 | 85.40 | 90.16 | 94.76 | 84.28 | 41.85 | 97.27 |
| Mean Temperature Warmest Quarter (˚C) | 147.34 | 155.41 | 161.40 | 167.15 | 165.30 | 119.03 | 161.93 |
| Minimum Annual Temperature (˚C) | 5.38 | 8.00 | 7.91 | 9.40 | 8.00 | 3.90 | 9.78 |
| Precipitation Driest Quarter (mm) | 123.47 | 53.08 | 154.69 | 131.51 | 125.20 | 300.53 | 153.06 |
| Precipitation Seasonality (mm) | 18.24 | 15.71 | 14.51 | 21.61 | 31.57 | 31.17 | 33.56 |
| Precipitation Wettest Quarter (mm) | 200.08 | 238.05 | 198.58 | 231.63 | 295.14 | 724.91 | 380.30 |
| Public Road Length (m) | 31415.93 | 263.30 | 918.77 | 243.95 | 98.45 | 307.94 | 1997.69 |
| Sealed Road Length (m) | 31415.93 | 263.31 | 61.65 | 420.78 | 671.02 | 245.23 | 53.83 |
| Surface Area of Bodies of Water ($m^2$) | 455.62 | 29627.00 | 124.44 | 16695.96 | 316.11 | 494.04 | 1133.77 |

**Supplementary Table 2**: Mean and standard error estimates for five of the abiotic environmental variables used in the landscape genomic analyses. Means were estimated by randomly sampling five points in the 25 $km^2$ ellipses produced for each sampling location. ± indicates standard error. There were no significant differences between the centroid values and the means for any environmental variable in any sampling locality.

| | **Isothermality** | **Mean Annual Temperature** (˚C) | **Precipitation seasonality (mm)** | **Elevation (m)** | **Vegetation Index** |
|---|---|---|---|---|---|
| **FEN** | 49.871 ± 0.07 | 10.4 ± 0.5 | 18.8 ± 0.7 | 257.4 ± 96.9 | 29.4 ± 17.9 |
| **FOR** | 51.40 ± 0.5 | 12 ± 0.2 | 16 ± 0 | 79.4 ± 24.8 | 59.6 ± 23.2 |
| **FRY** | 54.09 ± 0.03 | 12.5 ± 0.30 | 14.2 ± 0.40 | 105.2 ± 21.6 | 72.6 ± 5.9 |
| **MTW** | 49.88 ± 0.49 | 13.0 ± 0.1 | 22 ± 0.6 | 58.8 ± 26 | 46.4 ± 17.7 |
| **NAR** | 48.07 ± 0.04 | 12.5 ± 0.2 | 32 ± 0.6 | 68.8 ± 42.02 | 61.8 ± 11.9 |
| **WPP** | 4798 ± 0.01 | 7.9 ± 0.1 | 31.2 ± 0.4 | 713.4 ± 23.1 | 69.8 ± 4.2 |
| **WOO** | 49.92 ± 0.02 | 12.8 ± 0.04 | 33.4 ± 0.8 | 13.6 ± 4.9 | 39.4 ± 23.6 |

**Supplementary Table 3**: Mean Pi (π), Tajima's D and $F_{IS}$ values ± 95% confidence intervals calculated using putatively neutral SNPs for the five populations in which samples were collected both prior to and post-disease arrival.

| | π | | Tajima's D | | $F_{IS}$ | | $F_{ST}$ |
|---|---|---|---|---|---|---|---|
| | **Pre-DFTD** | **Post-DFTD** | **Pre-DFTD** | **Post-DFTD** | **Pre-DFTD** | **Post-DFTD** | **Pre VS Post-DFTD** |
| **FEN** | 0.339 ± 0.146 | 0.340 ± 0.101 | 1.032 ± 0.897 | 1.126 ± 0.879 | 0.3222 ± 0.346 | 0.314 ± 0.371 | 0.00014 ± 0.016 |
| **FOR** | 0.311 ± 0.166 | 0.318 ± 0.156 | 1.003 ± 0.912 | 1.167 ± 0.941 | 0.409 ± 0.389 | 0.430 ± 0.357 | 0.004 ± 0.017 |
| **FRY** | 0.332 ± 0.151 | 0.328 ± 0.149 | 1.243 ± 0.939 | 1.262 ± 0.936 | 0.258 ± 0.359 | 0.369 ± 0.355 | 0.0034 ± 0.008 |
| **NAR** | 0.342 ± 0.142 | 0.342 ± 0.143 | 1.229 ± 0.869 | 1.142 ± 0.865 | 0.256 ± 0.352 | 0.293 ± 0.364 | 0.002 ± 0.009 |
| **WPP** | 0.336 ± 0.153 | 0.339 ± 0.147 | 0.947 ± 0.879 | 1.251 ± 0.923 | 0.198 ± 0.389 | 0.204 ± 0.363 | 0.0001 ± 0.012 |

**Supplementary Table 4:** Summary of the PCA loading scores of the first six principal components for vegetative and geographic features for the seven sampling localities which cumulatively explain > 99% of the observed variance.

| | **PC 1** | **PC 2** | **PC 3** | **PC 4** | **PC 5** | **PC 6** |
|---|---|---|---|---|---|---|
| Eigenvalue (**$\lambda$**) | 8.48 | 4.98 | 2.11 | 1.59 | 0.70 | 0.13 |
| Cumulative proportion of variance explained | 0.471 | 0.277 | 0.117 | 0.088 | 0.039 | 0.007 |
| Mean Annual Temperature | 0.334 | 0.071 | -0.010 | 0.112 | 0.023 | 0.223 |
| Precipitation Seasonality | -0.135 | 0.240 | 0.149 | 0.558 | -0.080 | -0.283 |
| Isothermality | 0.192 | -0.062 | -0.237 | -0.427 | 0.598 | -0.256 |
| Annual Temperature Range | -0.097 | -0.353 | -0.296 | 0.144 | -0.319 | 0.277 |
| Vegetation Index | 0.021 | 0.287 | -0.517 | 0.034 | -0.169 | -0.062 |
| Length of Sealed Roads | -0.052 | -0.431 | 0.062 | 0.157 | 0.027 | -0.120 |
| Elevation | -0.339 | 0.005 | -0.007 | -0.127 | -0.018 | 0.106 |
| Surface Area of Water | 0.133 | 0.039 | 0.321 | -0.537 | -0.486 | 0.094 |

**Supplementary Table 5**: The 64 candidate genes detected pre-disease using landscape genomics analyses. ENSEMBL ID, NCBI gene function, NCBI gene ID, associated GO terms are included for each candidate gene. The seven candidate genes that were shared between the pre and post-disease candidate gene lists are colored grey.

| ENSEMBL ID | NCBI Function | Gene ID | GO Molecular Functions | GO Biological Processes | GO Cellular Components |
|---|---|---|---|---|---|
| ENSSHAG00000005439 | ankyrin repeat and MYND domain containing 2 | ANKMY2 | Metal ion binding; enzyme binding | - | Cilium; cell projection |
| ENSSHAG00000007940 | Armadillo repeat containing 3 | ARMC3 | - | - | - |
| ENSSHAG00000016653 | Actin related protein 2/3 complex subunit 2 | ARPC2 | Actin filament binding; structural molecule activity | Arp2/3 complex-mediated actin nucleation; actin filament polymerization | Actin cytoskeleton; cytoskeletal part; protein-containing complex |
| ENSSHAG00000011967 | BMP binding endothelial regulator | BMPER | - | Cell adhesion; reproduction | Extracellular matrix |
| ENSSHAG00000010409 | chromosome 4 open reading frame 17 | C4ORF17 | - | - | - |
| ENSSHAG00000011510 | Calnexin | CANX | Calcium ion binding | Exocytosis; intracellular protein transport; protein folding; protein metabolic process | - |
| ENSSHAG00000009956 | core-binding factor | CBFB | Sequence-specific DNA binding transcription coactivator activity | Nucleic acid-templated transcription | RNA polymerase II transcription factor complex |
| ENSSHAG00000007042 | Coiled-coil domain containing 162 | CCDC162 | - | - | - |
| ENSSHAG00000008512 | CD36 molecule | CD36 | - | Cell adhesion; cellular process | - |
| ENSSHAG00000007822 | centromere protein H | CENPH | Protein binding; kinetochore binding | Chromosome segregation; kinetochore assembly & organization; mitotic cell cycle | Kinetochore; nucleus; chromosome, centromeric region |
| ENSSHAG00000001719 | calmegin | CLGN | Calcium ion binding | exocytosis; protein metabolic process; intracellular protein transport; protein folding | - |
| ENSSHAG00000005741 | collagen | CO12A1 | - | - | - |
| ENSSHAG00000015846 | component of oligomeric golgi complex 1 | COG1 | - | - | Golgi apparatus; vacuole; plasma membrane |

| ENSEMBL ID | NCBI Function | Gene ID | GO Molecular Functions | GO Biological Processes | GO Cellular Components |
|---|---|---|---|---|---|
| ENSSHAG00000007801 | cut-like homeobox 1 | CUX1 | RNA polymerase-II activity; DNA binding; chromatin binding | Golgi vesicle transport; regulation of transcription by RNA polymerase II | Golgi membrane; nucleus; plasma membrane; vacuole |
| ENSSHAG00000016606 | cytochrome P450 | CYP39A1 | Iron ion binding; hydroxylase activity; oxidoreductase activity; heme binding | Lipid, bile acid, cholesterol and steroid metabolic process; digestion; oxidation-reduction process | Endoplasmic reticulum; membrane; intracellular membrane-bounded organelles |
| ENSSHAG00000015701 | elongator acetyltransferase complex subunit 4 | ELP4 | RNA polymearse II core binding | Regulation of transcrption by RNA polymerase II; transcription by RNA polymerase II | Cytoplasm; transcription elongation factor complex |
| ENSSHAG00000005180 | FGGY carbohydrate kinase domain containing [Source | FGGY | Transferase activity | Phosphate-containing compound metabolic process | Cytoplasm |
| ENSSHAG00000014922 | glucosidase | GBA3 | Glucosidase activity | - | - |
| ENSSHAG00000010474 | Uncharacterized protein | GM281 | Methylated histone binding | Anatomical structure morphogenesis; embryo development; spermatogenesis | Cytosol; synaptonemal complex; nucleoplasm |
| ENSSHAG00000015344 | glycosyltransferase-like domain containing 1 | GTDC1 | - | - | - |
| ENSSHAG00000003851 | major histocompatibility complex | HLA-DMA | Protein binding | Immune system process; immunoglobulin mediated immune response; response to stimulus | Lysosome; endosome |
| ENSSHAG00000008668 | Uncharacterized protein | KDM4C | Histone demethylase activity; oxidoreductase activity, incorporation of one atom each of oxygen into both donors | Chromatin remodeling | Histone methyltransferase complex |
| ENSSHAG00000004610 | kelch-like family member 20 | KLHL20 | Ubiquitin-protein transferase activity; actin binding | Proteasome-mediated ubiquitin-dependent protein catabolic process; golgi to endosome transport | Nucleus; cytoplasm; nuclear speck; golgi network; PML body; cell projection |
| ENSSHAG00000009141 | Kynureninase | KYNU | hydrolase activity | Aromatic amino acid family catabolism; cellular nitrogen compound catabolism; heterocycle catabolism | Cytoplasm |

| ENSEMBL ID | NCBI Function | Gene ID | GO Molecular Functions | GO Biological Processes | GO Cellular Components |
|---|---|---|---|---|---|
| ENSSHAG00000005591 | LIM domains containing 1 | LIMD1 | Transcription corepressor activity | Response to hypoxia; gene expression; gene silencing by miRNA; intracellular signal transduction & negative regulation; positive regulation of cellular process | P-body; adherens junction; cell-cell junction; nucleus; transcription factor complex |
| ENSSHAG00000003547 | mediator of cell motility 1 | MEMO1 | - | Cellular process; regulation of biological process | Cytoplasm |
| ENSSHAG00000007088 | Uncharacterized protein | MFRP | - | Chemical synaptic transmission; sensory perception | - |
| ENSSHAG00000009275 | Alpha-1,3-mannosyl-glycoprotein 4-beta-N-acetylglucosaminyltransferase A | MGAT4A | Metal ion binding; transferase activity; transferring glycosol groups | Protein N-link glycosylation | Golgi apparatus; integral component of membrane; extracellular region |
| ENSSHAG00000013191 | alpha-1,6-mannosylglycoprotein 6-beta-N-acetylglucosaminyltransferase | MGAT5 | Acetylglucosaminyltransferase activity; catalytic activity, acting on a protein | Protein N-linked glycosylation | Golgi apparatus; vacuole; plasma membrane |
| ENSSHAG00000016108 | myeloid/lymphoid or mixed-lineage leukemia (trithorax homolog | MLLT3 | DNA-binding transcription factor activity; chromatin binding | Transcription by RNA polymerase II | - |
| ENSSHAG00000014273 | membrane protein | MPP2 | Kinase activity | - | - |
| ENSSHAG00000013782 | mitochondrial ribosomal protein L43 | MRPL43 | Structural constituent of ribosome | - | Mitochondrial large ribosomal subunit |
| ENSSHAG00000000563 | Myeloid-derived growth factor | MYDGF | - | Negative regulation of apoptotic process; postivie regulation of endothelial cell proliferation; positive regulation of MAPK cascade; postive regulation of angiogenesis | Extracellular matrix; endoplasmic reticulum; golgi apparatus |
| ENSSHAG00000000752 | Myosin IC | MYO1C | Enzyme regulator activity; motor activity; protein binding | Cytokinesis; intracellular protein transport & signal transduction; sensory perception of sound; muscle contraction | actin cytoskeleton; plasma membrane |

| ENSEMBL ID | NCBI Function | Gene ID | GO Molecular Functions | GO Biological Processes | GO Cellular Components |
|---|---|---|---|---|---|
| ENSSHAG00000010248 | natriuretic peptide receptor 1 | NPR1 | Adenylate cyclase activity; peptide receptor activity | Cyclic nucleotide metabolic process; purine ribonucleotide biosynthetic process; enzyme linked receptor protein signaling pathway | - |
| ENSSHAG00000000012 | Olfacorty Receptor 52L2 | OR52L1 | - | - | - |
| ENSSHAG00000006602 | peptidase domain containing associated with muscle regeneration 1 | PAMR1 | Peptidase activity | Response to external stimulus | - |
| ENSSHAG00000013654 | PDZ domain containing 7 | PDZD7 | - | - | Plasma membrane |
| ENSSHAG00000013620 | Protein phosphatase 1 regulatory subunit 12B | PPP1R12B | Enzyme inhibitor activity; phosphatase activity & regulation | - | A band; Z disc |
| ENSSHAG00000012415 | presenilin 1 | PSEN1 | Endopeptidase activity | Calcium ion transport; cellular protein & peptide metabolic process; membrane protein proteolysis | Z disc; apical plasma membrane; cell cortex; perinuclear region of cytoplasm; nucleus; neuronal cell body |
| ENSSHAG00000014446 | receptor (TNFRSF)-interacting serine-threonine kinase 1 | RIPK1 | JUN kinase activity; MAP kinase activity | Activation of JUN kinase activity & MAPK activity | Intracellular |
| ENSSHAG00000011219 | ribosomal protein L7-like 1 | RPL7L1 | RNA binding; structural constituent of ribosome | Maturation of LSU-rRNA from tricistronic rRNA transcript (SSU-rRNA, 5.8S rRNA, LSU-rRNA) | Cytosolic large ribosomal subunit |
| ENSSHAG00000018275 | ribosomal RNA processing 12 homolog | RRP12 | - | - | Integral component of membrane |
| ENSSHAG00000001518 | Sodium voltage-gated channel alpha subunit 9 | SCN9A | Cation channel activity; sodium ion transmembrane transporter activity | Action potential; chemical synaptic transmission; nervous system process | Cation channel complex; integral component of plasma membrane |
| ENSSHAG00000015982 | surfactant protein B | SFTPB | - | - | Extracellular space; late endosome; plasma membrane; secretory granuale; vacuole |
| ENSSHAG00000013892 | SH3-domain kinase binding protein 1 | SH3KBP1 | SH3 domain binding; ubiquitin protein ligase binding | Cell migration; regulation of cell shape; apoptotic process; postivie regulation of epidermal growth factor receptor signaling | Cytoplasm; endocytic vesicle; cell junction; neuron projection |

| ENSEMBL ID | NCBI Function | Gene ID | GO Molecular Functions | GO Biological Processes | GO Cellular Components |
|---|---|---|---|---|---|
| ENSSHAG00000016788 | solute carrier family 39 | SLC39A11 | Metal ion transmembrane transporter activity | Divalent metal ion transport; inorganic cation transmembrane transport; transition metal ion transport | - |
| ENSSHAG00000008676 | solute carrier family 4 (sodium bicarbonate cotransporter) | SLC4A5 | Organic anion transmembrane transporter activity; sodium ion transmembrane transporter Activity; solute:cation symporter activity | - | - |
| ENSSHAG00000007350 | solute carrier organic anion transporter family | SLCO4C1 | Organic anion transmembrane transporter activity; secondary active transmembrane transporter activity | - | Integral component of plasma membrane |
| ENSSHAG00000012499 | syntaxin 17 | STX17 | SNARE binding | Intracellular protein transport; organelle localizaiton by membrane tethering; vesicle fusion | SNARE complex; endomembrane system; integral component of membrane; plasma membrane; vacuole |
| ENSSHAG00000014392 | syntaxin 2 | STX2 | SNARE binding | Vesicle fusion to plasma membrane; intracellular protein transport; organelle localization by membrane tethering; synaptic vesicle exocytosis | SNARE complex; synaptic vesicle; vacuole; presynapic active zone; ingegral component of membrane; presynaptic membrane |
| ENSSHAG00000009436 | synaptotagmin I | SYT1 | Calcium-dependent phospholipid binding; phosphatidylserine binding; syntaxin binding | Organic hydroxy compound transport; cellular response to stimulus; ammonium transport; drug transport; synaptic vesicle endocytosis & exocytosis; vesicle budding & fusion to membrane; regulation of exocytosis & ion transport | Axon; vacuole; synaptic vesicle membrane; plasma membrane region; secretory granule |
| ENSSHAG00000013165 | TBC1 | TBC1D1 | GTPase activator activity; GTPase binding | Intracellular protein transport | Intracellular |
| ENSSHAG00000013042 | Thromboxane A synthase 1 | TBXAS1 | Oxidoreductase activity; isomerase activity; heme binding; metal ion binding | Fatty acid & lipid metabolism; oxidation-reduction process; cellular chloride ion homeostasis; positive regulation of vasoconstriction | Integral component of membrane; endoplasmic reticulum |

| ENSEMBL ID | NCBI Function | Gene ID | GO Molecular Functions | GO Biological Processes | GO Cellular Components |
|---|---|---|---|---|---|
| ENSSHAG00000013677 | transmembrane and coiled-coil domain family 2 | TMCC2 | - | - | Endomembrane system; vacuole; plasma membrane |
| ENSSHAG00000007483 | transmembrane protein 181 | TMEM181 | Toxic substance binding; pathogenesis | - | - |
| ENSSHAG00000000473 | transmembrane protein 231 | TMEM231 | - | Protein localization & regulation of protein localization | Bounding membrane of organelle; intraciliary transport particle; ciliary transition zone; cell projection membrane |
| ENSSHAG00000013176 | transmembrane protein 68 | TMEM68 | Transferase activity, transferring acyl groups | - | Integral component of plasma membrane |
| ENSSHAG00000017996 | Tumor Necrosis Factor receptor superfamily member 21 | TNFRSF21 | - | B cell & T cell proliferation; bleb assembly; apoptotic signaling; adaptive immune response; mylineation; humoral immune response | Integral component of membrane |
| ENSSHAG00000000170 | tripartite motif containing 10 | TRIM10 | Zinc ion binding; metal ion binding | Innate immune responses; erythrocyte differentiation; negative regulation of viral entry into host cell | Cytoplasm |
| ENSSHAG00000004397 | Uncharacterized protein | UNK | - | - | - |
| ENSSHAG00000013089 | ubiquitin specific peptidase 42 | USP42 | Cysteine-type endopeptidase activity; thiol-dependent ubiquitin-specific protease activity | Apoptotic signaling pathway; bleb assembly; execution & regulation of apoptosis; protein deubiquitination | - |
| ENSSHAG00000017298 | KIAA1429 | VIRMA | - | mRNA processing, alternative polyadenylation & processing; RNA splicing; multicellular organism development | Nucleus; cytoplasm; nuclear speck; cytosol; RNA N6-methyladenosine methyltransferase complex |
| ENSSHAG00000014544 | zinc finger | ZCCHC7 | Zinc ion binding; nucleic acid binding | - | Cytosol, nucleus, nucleolus |

**Supplementary Table 6**: The 76 candidate genes uniquely detected post-disease using landscape genomics analyses. Each candidate gene is identifiable with an ENSEMBL ID, and an NCBI gene function and gene ID, and includes all of the associated GO terms.

| ENSEMBL ID | NCBI Function | Gene ID | GO Molecular Functions | GO Biological Processes | GO Cellular Components |
|---|---|---|---|---|---|
| ENSSHAG00000012498 | activin A receptor | ACVR2B | Activin binding; cytokine receptor binding; growth factor binding | Activin receptor signaling pathway; transforming growth factor beta receptor signaling pathway | Integral component of plasma membrane; serine/threonine protein kinase complex |
| ENSSHAG00000018155 | ADAM metallopeptidase with thrombospondin type 1 motif | ADAMTS18 | Peptidase inhibitor activity; protein binding; metallopeptidase activity | Cellular process | Extracellular region |
| ENSSHAG00000014822 | ADAM metallopeptidase with thrombospondin type 1 motif | ADAMTS9 | Metallopeptidase & endopeptidase activity | Heat valve morphogenesis; proteolysis; response to bacteria; melanoycte differentiation | Endoplasmic reticulum; extracellular matrix; cell surface |
| ENSSHAG00000005258 | Alcohol dehydrogenase 1-lik | ADH6A | Oxidoreductase activity; zinc ion binding | Alcohol metabolic process; cellular response to oxygen-containing compounds; responses to toxic substances; hormone metabolism | Cytosol |
| ENSSHAG00000015563 | Rho guanine nucleotide exchange factor (GEF) 37 | ARHGEF37 | GTP binding; guanyl-nucleotide exchange factor activity | Regulation of Rho protein signal transduction | Cytoplasm |
| ENSSHAG00000004447 | ADP-ribosylation factor-like 8B | ARL8B | Nucleotide binding; GRP binding; alpha-tubulin binding | Cell division; lysosome localization; anterograde axonal transport | Cytoplsm; cytoskeleton; lysosome; synapse |
| ENSSHAG00000000974 | aspartate beta-hydroxylase | ASPH | Calcium ion binding; dioxygenase activity; oxidoreductase activity | Negative regulation of cell population proliferation; response to ATP; pattern specification process; regulation of protein stability | Endoplasmic reticulum; Cytoplasm; integral component of membrane |
| ENSSHAG00000011416 | BicC family RNA binding protein 1 | BICC1 | RNA-binding; lipid transporter activity | Cell cycle; developmental process; lipid metabolic process | Cytoplasm |
| ENSSHAG00000011967 | BMP binding endothelial regulator | BMPER | Extracellular matrix structural constituent | Regulation of endothelial cell migration; regulation of angiogenesis | Extracellular space |
| ENSSHAG00000003636 | chromosome 8 open reading frame 37 | C8ORF37 | Protein binding | Photoreceptor cell morphogenesis | Cytoplasm |

| ENSEMBL ID | NCBI Function | Gene ID | GO Molecular Functions | GO Biological Processes | GO Cellular Components |
|---|---|---|---|---|---|
| ENSSHAG00000004855 | calbindin 1 | CALB1 | Calcium ion binding | Anatomical structure homeostasis; chemical synaptic transmission; regulation of cytosolic calcium ion concentration | Cytosol; dendrite; nucleus; plasma membrane region; terminal bouton |
| ENSSHAG00000011323 | CD8b molecule | CD8B | Protein binding | Adaptive immune response; immune system process | External side of T cell of plasma membrane |
| ENSSHAG00000017154 | cadherin 8 | CDH8 | cadherin binding; protein homodimerization activity; calcium ion binding; cytoskeletal protein binding | Adherens junction organization; cell-cell junction assembly | Cell-cell adherens junction; plasma membrane protein complex |
| ENSSHAG00000002539 | cyclin-dependent kinase 19 | CDK19 | Cyclin-dependent protein serine/threonine kinase activity; molecular transducer activity | Cell cycle | Core mediator complex; mediator complex |
| ENSSHAG00000010121 | contactin 3 (plasmacytoma associated) | CNTN3 | - | Cell adhesion; nervous system development | Neuron projection; anchored component of plasma membrane |
| ENSSHAG00000004917 | collagen | COL13A1 | Extracellular matrix structural constituent | Extracellular matrix organization | Extracellular matrix |
| ENSSHAG00000014768 | collagen | COL27A1B | - | - | Extracellular space |
| ENSSHAG00000014932 | cytoplasmic polyadenylation element binding protein 4 | CPEB4 | mRNA 3'-UTR binding; translation regulator activity | Negative regulation of translation; formation of translation initiation ternary complex | Cytoplasm; neuron projection; ribonucleoprotein complex |
| ENSSHAG00000017544 | cartilage acidic protein 1 | CRTAC1 | Calcium ion binding | Axonal fasciculation; olfactory bulb development | Growth cone; extracellular region |
| ENSSHAG00000003551 | CUB and Sushi multiple domains 2 | CSMD2 | - | - | - |
| ENSSHAG00000014221 | diaphanous-related formin 3 | DIAPH3 | Actin binding; Rho GTPase binding | Movement of cell or subcellular component | Actin cytoskeleton |
| ENSSHAG00000011075 | dispatched homolog 1 (Drosophila) | DISP1 | Peptide transmembrane transporter activity | Cell surface receptor signaling pathway | Integral component of membrane |
| ENSSHAG00000007803 | deleted in malignant brain tumors 1 | DMBT1 | Scavenger receptor activity; extracellular matrix structural constituent; zymogen binding | Blastocyst development; response to bacterium; mass & epithelial cell proliferation | External side of plasma membrane; leaflet of membrane bilayer |
| ENSSHAG00000009824 | DnaJ (Hsp40) homolog | DNAJA3 | Protein kinase binding; Hsp70 protein binding | Apoptotic process; activation-induced death of T cells; cell aging; heat response | Nucleus; Cytoplasm; mitochondrion |

| ENSEMBL ID | NCBI Function | Gene ID | GO Molecular Functions | GO Biological Processes | GO Cellular Components |
|---|---|---|---|---|---|
| ENSSHAG00000002033 | dedicator of cytokinesis 5 | DOCK5 | GTPase activity | Intracellular signal transduction | Cytosol |
| ENSSHAG00000014792 | epilepsy | EMP2A | - | - | - |
| ENSSHAG00000017567 | Folliculin | FLCN | Protein-containing complex binding; guanyl-nucleotide exchange factor activity | Negative regulation of transcription of RNA polymerase II; negative regulation of angiogenesis; regulation of histone acetylation | Cell-cell contact zone; midbody |
| ENSSHAG00000016322 | putative dimethylaniline monooxygenase [N-oxide-forming] 6 | FMO6 | Monooxygenase activity | - | - |
| ENSSHAG00000010151 | follistatin-like 1 | FSTL1 | Heparin binding; calcium ion binding | Response to starvation | Extracellular space |
| ENSSHAG00000011468 | glutamate decarboxylase 1 (brain | GAD1 | Lyase activity | Cellular amino acid metabolic process | Cytoplasm; axon; mitochondrion |
| ENSSHAG00000012829 | growth hormone receptor | GHR | Cytokine receptor activity; growth factor binding; peptide hormone binding | JAK-STAT cascade; cellular response to peptide hormone stimulus; peptidyl-tyrosine phosphorylation | Cytosol; receptor complex; leaflet of membrane bilayer |
| ENSSHAG00000014713 | GTP binding protein 4 | GTPBP4 | GTPase activity; GTP & nucleotide binding | Negative regulation of DNA replication; negative regulation of cell population;ribosome biogenesis proliferationstranslation | Cytoplasm; nucleus; golgi apparatus |
| ENSSHAG00000002512 | histone deacetylase 7 | HDAC7 | Deacetylase activity; nucleic acid binding; oxidoreductase activity | Negative regulation of apoptotic process; chromatin organization | Histone deacetylase complex; nucleus; cytosol |
| ENSSHAG00000009284 | HtrA serine peptidase 1 | HTRA1 | Peptidase activity | Intracellular signal transduction; protein folding | Cytoplasm; plasma membrane; extracellular space |
| ENSSHAG00000002618 | interferon-induced protein with tetratricopeptide repeats 5-like | IFIT5 | RNA binding | Defense response to virus | Cytosol |
| ENSSHAG00000009342 | interleukin 9 receptor | IL9R | Cytokine receptor activity; interleukin-9 receptor activity | Cellular process; response to stimulus; cellular inflamamatory response | Integral component of plasma membrane |
| ENSSHAG00000008101 | IQ motif and ubiquitin domain containing | IQUB | Protein binding | Smoothened signaling pathway; cilium assembly; cell projection organization | Motile cilium; acrosomal vesicle |
| ENSSHAG00000010089 | integrin alpha FG-GAP repeat containing 1 | ITFG1 | Protein binding | Modulator of T-cell function; response to stimulus | Integral component of plasma membrane |

| ENSEMBL ID | NCBI Function | Gene ID | GO Molecular Functions | GO Biological Processes | GO Cellular Components |
|---|---|---|---|---|---|
| ENSSHAG00000003020 | kazrin | KAZN | - | Keratinization | Cornified envelope; nuclear speck; desmosome |
| ENSSHAG00000004610 | Kelch like family member 20 | KLHL20 | Actin binding; ubiquitin-protein transferase activity; interferon-gamma binding | Golgi to endosome transport; regualtion of apoptotic process; protein ubiquitination | PML body; cytosol; golgi apparatus |
| ENSSHAG00000011783 | monoglyceride lipase | MGLL | Hydrolase activity; acylglycerol lipase activity; protein homodizeration activity; carboxylic ester hydrolase activity | Lipid metabolic process; regulation of signal transduction; regulation of inflammatory response & sensory perception of pain | Cytoplasm; synapse; cytosol; varcosity |
| ENSSHAG00000001599 | methylmalonyl CoA mutase | MMUT | GTPase activity; protein binding; catalytic activity | Positive regulation of GTPase activity; metabolic process | Mitochondrion |
| ENSSHAG00000002966 | 5-methyltetrahydrofolate-homocysteine methyltransferase | MTR | Cellular amino acid biosynthetic process; sulfur compound metabolic process | RNA processing; RNA catabolism and splicing; translation termination; DNA methylation | Exosome (Rnase complex); TRAMP; nucleus; cytosol |
| ENSSHAG00000015937 | myosin XVIIIA | MYO18A | - | Anatomical structure morphogenesis; muscle contraction; sensory perception of sound | - |
| ENSSHAG00000018189 | myosin ID | MYO1D | Enzyme regulator activity; structural molecule activity; motor activity | Cellular component morphogenesis; sensory perception of sound; muscle contraction; cytokinesis | Actin cytoskeleton; cell junction; plasma membrane |
| ENSSHAG00000017586 | NADH dehydrogenase (ubiquinone) Fe-S protein 3 | NDUFS3 | NADH dehydrogenase activity; oxidoreducate activity, activity on NAD(P)H | Oxidation-reduction process; reactive oxygen species metabolism; negative regulation of cell growth negative regulation of intrainsc apoptotic signaling pathway | ER to Golgi transport vesicle membrane |
| ENSSHAG00000017306 | neogenin 1 | NEO1 | Protein binding; signaling receptor activity; adherin binding; BMP receptor binding | Neuron migration; regulation of transcription, DNA-templated; cell adhesion; axon regeneration | Nucleoplasm; golgi apppartaus; cell surface |
| ENSSHAG00000010452 | olfactory receptor 51E2 | OR51E2 | Transmembrane signling receptor activity | Detection of chemical stimulus involved in sensory perception | Plasma membrane |
| ENSSHAG00000018561 | oxidation resistance 1 | OXR1 | Oxidoreducatse activity | Neuron apoptotic process; cellular response to hydroperoxidase | Nucleus; mitochondrion; nucleoplasm |
| ENSSHAG00000007227 | pericentriolar material 1 | PCM1 | Protein binding in centrosome | Intraciliary transport involved in cilium assembly; centrosome cycle | Centrosome; intraciliary transport particle; intraciliary transport particle |

| ENSEMBL ID | NCBI Function | Gene ID | GO Molecular Functions | GO Biological Processes | GO Cellular Components |
|---|---|---|---|---|---|
| ENSSHAG00000010292 | pleckstrin homology domain containing | PLEKHA3 | Amide binding; phospholipid binding& transporter activity | Amide transport; membrane organization | Cytosol |
| ENSSHAG00000014793 | perilipin 2 | PLIN2 | Protein binding | Lipid metabolic process | Part of trophoblast cell; lipid droplet; nucleus |
| ENSSHAG00000013620 | protein phosphatase 1 regulatory subunit 12B | PPP1R12B | Enzyme inhibitor activity; phosphatase activity | Signal transduction; regulates myosin phosphatase | A band; Z disc |
| ENSSHAG00000009819 | PR domain containing 11 | PRDM11 | Chromatin DNA binding; RNA polymerase II regulatory region sequence-specific DNA binding | Positive regulation of transcription, DNA-templated | Nucleus |
| ENSSHAG00000017902 | protein tyrosine phosphatase | PTPRG | Phosphoprotein phosphatase activity | Extracellular matrix organization | Cytoplasm |
| ENSSHAG00000006991 | protein tyrosine phosphatase | PTPRK | Phosphorpotein phosphatase activity | Protein dephosphorylation; cellular response to UV & reactive oxygen species; negative regulation of cell population proliferation | Cytoplasm |
| ENSSHAG00000008814 | RING FINGER PROTEIN 126 | RFN126 | E3 ligase | Ubiquination of AID (Activation-induced cytidine deaminase) in immunoglobulin genes | - |
| ENSSHAG00000007644 | RH-like protein | RHD | Ammonium transmembrane transporter activity | - | Integral component of plasma membrane |
| ENSSHAG00000016431 | rabphilin 3A | RPH3A | Zinc ion, protein & lipid binding | Regulation of exocytosis; glucose homeostasis; regulation of protein secretion; spontaneous neurotransmitter secretion | Cytosol; synaptic vesicle; cell junction |
| ENSSHAG00000014744 | sodium channel and clathrin linker 1 | SCLT1 | Protein C-terminus binding; sodium channgel regulator activity; clathrin binding | Cilium assembly; clustering of voltage gated sodium channels | Centrosome; ciliary transport; cytoskeleton |
| ENSSHAG00000015424 | SEC31 homolog B (S. cerevisiae) | SEC31B | Structural molecule activity | COPII-coated vesicle budding; endoplasmic reticulum organization | ER to Golgi transport vesicle membrane; endoplasmic reticulum exit site |
| ENSSHAG00000003571 | solute carrier family 15 (oligopeptide transporter) | SLC15A1 | Dipeptide transmembrane transporter activity; solute:proton symporter activity | Oligopeptide & peptide transport; negative regulation of amino acid transport | Integral component of plasma membrane |

| ENSEMBL ID | NCBI Function | Gene ID | GO Molecular Functions | GO Biological Processes | GO Cellular Components |
|---|---|---|---|---|---|
| ENSSHAG00000001366 | solute carrier family 26 (anion exchanger) | SLC26A11 | Methyltransferase activity | In the same pathway as ASPH | Integral component of plasma membrane |
| ENSSHAG00000008623 | Sodium-coupled neutral amino acid transporter 1 | SLC38A1 | Amino acid transmembrane transporter activity | - | Integral component of plasma membrane |
| ENSSHAG00000005902 | solute carrier family 7 (cationic amino acid transporter | SLC7A1 | L-lysine transmembrane transporter activity; arginine transmembrane transporter activity | Import into cell | Cytoplasm; apical dendrite; lysosome |
| ENSSHAG00000014163 | SMG5 nonsense mediated mRNA decay factor | SMG5 | RNA-directed DNA polymerase activity; telomeric DNA binding | Regulation of DNA metabolic process; telomere maintenance & assembly; telomere maintenance & assembly | Cytoplasm; nuclear part; ribonucleoprotein complex |
| ENSSHAG00000004975 | chromosome 9 open reading frame 9 | SPACA9 | - | Acrosomal vesicle; ciliary transition; axonemal central apparatus | |
| ENSSHAG00000014837 | syntaxin 12 | STX12 | SNARE binding | Intracellular protein transport; vesicle fusion | SNARE complex; integral component of membrane |
| ENSSHAG00000018351 | SUN domain containing ossification factor | SUCO | - | Ossification; regulation of collagen biosynthetic process & osteoblast differentiation | Endoplasmic reticulum; Cytoplasm; integral component of membrane |
| ENSSHAG00000007292 | syntabulin (syntaxin-interacting) | SYBU | Kineasin binding; syntaxin-1 binding | Regulation of synaptic activity; regulation of insulin secretion; axonal transport of mitochondrion | Golgi apparaturs; microtubule; axon |
| ENSSHAG00000017344 | transcription factor 7-like 2 (T-cell specific | TCF7L2 | Protein binding; sequence specific binding; transcription bidning | Canonical Wnt signaling pathway; regulation of transcription; insulin secretion; muscle structure development; cell fate commitment; positive regulation of cell communication | Nucleus; transcription factor complex |
| ENSSHAG00000017708 | thymidine kinase 2 | TK2 | Kinase activity | Nucleoside monophosphate biosynthetic process; cell cycle regulation | Cytoplasm |
| ENSSHAG00000010462 | Toll-like receptor | TLR6 | Transmembrane signling receptor activity; toll-like receptor 2 binding; lipopeptide binding | Innate immune response; inflammatory response; toll-like receptor signaling pathway | Integral component of plasma membrane |
| ENSSHAG00000014783 | trafficking protein | TRAK1 | Cytoskeletal protein binding; signaling receptor binding | Axo-dendritic transport; mitochondrion organization; neurogensis; protein targeting | SNARE complex; integral component of membrane |

| ENSEMBL ID | NCBI Function | Gene ID | GO Molecular Functions | GO Biological Processes | GO Cellular Components |
|---|---|---|---|---|---|
| ENSSHAG00000004802 | transient receptor potential cation channel subfamily V member 6 | TRPV6 | Calcium channel activity | Transport & response to calcium ions; parathyroid hormone secretion; | Integral component of plasma membrane |
| ENSSHAG00000015818 | Uncharacterized protein | UNK | - | - | - |
| ENSSHAG00000002503 | gastrula zinc finger protein XlCGF26.1-like | UNK | mRNA binding | Cell morphogenesis involved in neuron differentiation | - |
| ENSSHAG00000000903 | olfactory receptor 2AG1-like | UNK | - | - | - |
| ENSSHAG00000018392 | Uncharacterized protein | UNK | - | - | - |
| ENSSHAG00000001909 | Uncharacterized protein | UNK | - | - | - |
| ENSSHAG00000006847 | WD repeat domain 27 | WDR27 | - | - | Nucleoplasm |
| ENSSHAG00000015540 | xylosyltransferase I | XYLT1 | Transferase activity; metal ion binding | Ossification involved in bone maturation; cellular response to heat; negative regulation of axon regeneration | Golgi membrane; extracellular space |
| ENSSHAG00000003741 | zinc finger CCCH-type containing 18 | ZC3H18 | Metal ion binding | - | Nuclear speck; nucleus |

**Supplementary Table 7:** The 57 candidate genes detected uniquely pre-disease and the abiotic environmental variable(s) which the genetic-environmental association analyses detected statistically significant associations.

| Gene ID | Mean Annual Temperature | Seasonal Precipitation | Isothermality | Annual Temperature Range | Length of Sealed Roads | Elevation | Surface Area of Bodies of Water | Vegetation Index |
|---|---|---|---|---|---|---|---|---|
| ANKMY2 | X | | | | | | | |
| BMPER | X | | | | | | | |
| C4ORF17 | | | | | X | | X | |
| CBFB | | | X | | | | | X |
| CCDC162 | X | | | | | | | |
| CD36 | | | | | X | | | |
| CENPH | X | | | | | | | |
| CLGN | X | | X | | | | | X |
| CO12A1 | | | | X | | | | X |
| COG1 | | | | | X | | | |
| CUX1 | | | X | | | | | X |
| CYP39A1 | | X | | | | | X | |
| ELP4 | | | X | | | | | X |
| GBA3 | X | | | X | | | | |
| GM281 | | | | | X | | | |
| GTDC1 | X | | | | | | | |
| HLA-DMA | | | X | X | | | | X |
| KDM4C | | | | | | X | | |
| KLHL20 | | X | | | | X | X | |
| KYNU | | X | | | | | X | |
| LIMD1 | | | | X | | | | |
| MEMO1 | X | | | | | | | |
| MFRP | X | | | | | | | |
| MGAT5 | | | | | | X | | |
| MLLT3 | | X | | X | | | X | |
| MPP2 | | | | X | | | | |
| MRPL43 | | | X | | | | | X |

| Gene ID | Mean Annual Temperature | Seasonal Precipitation | Isothermality | Annual Temperature Range | Length of Sealed Roads | Elevation | Surface Area of Bodies of Water | Vegetation Index |
|---|---|---|---|---|---|---|---|---|
| MYDGF | | | | X | | | | |
| MYO1C | | | | | X | | | |
| NPR1 | X | | | | | | | |
| OR52L1 | X | | | | | | | |
| PAMR1 | X | | | | | | | |
| PDZD7 | | | | | X | X | | |
| PPP1R12B | | X | | | X | | | X |
| PSEN1 | | | X | | | | | X |
| RIPK1 | | | | | X | X | | |
| RPL7L1 | | | X | | | | | X |
| RRP12 | | X | | | | | X | |
| SFTPB | | | X | | | | | X |
| SH3KBP1 | | | | | | | X | |
| SLC39A11 | | | | | | X | | |
| SLC4A5 | | X | | | | | X | |
| SLCO4C1 | X | | | | | X | | |
| STX17 | X | | | | | | | |
| STX2 | | | X | | | | | X |
| SYT1 | X | | | | | | | |
| TBC1D1 | | X | | | | | X | |
| TMCC2 | | | | | X | | | |
| TMEM181 | | | | | | | X | |
| TMEM231 | | | | X | | | | |
| TMEM68 | | X | | | | | X | |
| TNFRSF21 | | X | | | | | | |
| TRIM10 | X | | | | | | | |
| UNK | | | X | | | | | X |
| USP42 | X | | | | | | | |
| VIRMA | | | | X | | | | |
| ZCCHC7 | | X | | | | | | |

**Supplementary Table 8:** The 76 candidate genes uniquely detected post-disease and the abiotic and biotic environmental variable(s) which the genetic-environmental association analyses detected statistically significant associations.

| Gene ID | Mean Annual Temperature | Seasonal Precipitation | Isothermality | Annual Temperature Range | Length of Sealed Roads | Elevation | Surface Area of Bodies of Water | Vegetation Index | Disease Prevalence |
|---|---|---|---|---|---|---|---|---|---|
| ACVR2B | | | | | | | | X | |
| ADAMTS18 | X | | | | | | | | |
| ADAMTS9 | | | | | | X | | | X |
| ADH6A | X | | | | | | | | |
| ARHGEF37 | | | | X | | | | | X |
| ARL8B | | | | X | | | | | |
| ASPH | | | | X | | | | | |
| BICC1 | | | | | | | | X | |
| BMPER | | X | | | | | | | |
| C8ORF37 | | X | X | | | | | | |
| CALB1 | | | X | | | | | | |
| CD8B | | | | | | X | | | |
| CDH8 | | | | | | | X | | |
| CDK19 | | | X | | | | | | |
| CNTN3 | | | | | | | | X | |
| COL13A1 | X | | | | X | | | | |
| COL27A1B | | | | | | | | X | |
| CPEB4 | | X | | | | | X | | |
| CRTAC1 | | | | | | | X | X | |
| CSMD2 | | | | | X | X | | | |
| DIAPH3 | | | | | | | | | X |
| DISP1 | | X | | | X | X | X | | |
| DMBT1 | | X | X | X | | | | X | X |

| Gene ID | Mean Annual Temperature | Seasonal Precipitation | Isothermality | Annual Temperature Range | Length of Sealed Roads | Elevation | Surface Area of Bodies of Water | Vegetation Index | Disease Prevalence |
|---|---|---|---|---|---|---|---|---|---|
| DNAJA3 | X | X | | | | | X | | |
| DOCK5 | | | | X | | | | | |
| EMP2A | | X | | | X | X | X | | |
| FLCN | | | | | | | | | X |
| FMO6 | | | | | X | | | | |
| FSTL1 | X | X | | | | | | | |
| GAD1 | | | X | | | | | | |
| GHR | | | X | | | | | | |
| GTPBP4 | | | X | | | | | | |
| HDAC7 | X | | | | | | | | |
| HTRA1 | | | X | | | | | | |
| IFIT5 | | | | X | | | | | |
| IL9R | | | | | | | | | X |
| IQUB | | | | X | | | | | |
| ITFG1 | | | | X | | | | | X |
| KAZN | X | | | | | | | | |
| KLHL20 | X | | | X | | | | | |
| MGLL | | | | | X | | | | |
| MMUT | | | | | | | | X | |
| MTR | | | | X | | | | | |
| MYO18A | | | | | | | | X | |
| MYO1D | | | | | | X | | | |
| NDUFS3 | | | | | | | | X | |
| NEO1 | X | | | | | | | | |
| OR51E2 | | | | | X | | X | | |
| OXR1 | | | | X | | | | | |
| PCM1 | | | | | | | | X | |

| **Gene ID** | **Mean Annual Temperature** | **Seasonal Precipitation** | **Isothermality** | **Annual Temperature Range** | **Length of Sealed Roads** | **Elevation** | **Surface Area of Bodies of Water** | **Vegetation Index** | **Disease Prevalence** |
|---|---|---|---|---|---|---|---|---|---|
| PLEKHA3 | | | | | X | X | | | |
| PLIN2 | | | | | | | | X | |
| PPP1R12B | | | | | | | | | X |
| PRDM11 | X | | | | | | | | |
| PTPRG | | | X | | | | | | |
| PTPRK | | | X | | | | | | |
| RFN126 | | | | | | | | | X |
| RHD | | | X | | | | | | |
| RPH3A | | | | | | | X | | |
| SCLT1 | | | | | | X | | | |
| SEC31B | | | X | | | | | | |
| SLC15A1 | | | | X | | | | | |
| SLC26A11 | | | | | | | | X | |
| SLC38A1 | | X | | | | X | X | | |
| SLC7A1 | | | | | | | | X | |
| SMG5 | | | | X | | | | | |
| SPACA9 | | | | | | | X | | |
| STX12 | | X | X | | | | | | |
| SUCO | | | | | | | | X | |
| SYBU | | | | | | | | | X |
| TCF7L2 | | | | | | | | X | |
| TK2 | | X | | | | | | | X |
| TLR6 | X | | | | | | | | |
| TRAK1 | | | | X | | | | | |
| TRPV6 | | | | | X | | | | |
| UNK | | | | | | | | X | |
| UNK | | | | | | | | | X |

| Gene ID | Mean Annual Temperature | Seasonal Precipitation | Isothermality | Annual Temperature Range | Length of Sealed Roads | Elevation | Surface Area of Bodies of Water | Vegetation Index | Disease Prevalence |
|---|---|---|---|---|---|---|---|---|---|
| UNK | | X | | | | | | | |
| UNK | | | | | | X | | | |
| UNK | X | | | X | | | X | | |
| WDR27 | | | | | X | X | | | |
| XYLT1 | X | | | | | | | | |
| ZC3H18 | | | | X | | | | | |

**Supplementary Table 9:** The seven candidate genes that were detected with significant genetic-environmental associations both pre- and post-disease and the abiotic/biotic environmental variable(s) which had statistically significant associations. Genetic-environmental associations are indicated with X for pre-disease (blue) and post-disease (orange) associations. Blue and orange crosses in the same cell reflect that the genetic-environmental association was detected both prior to and post-disease arrival.

| Gene ID | Mean Annual Temperature | Seasonal Precipitation | Isothermality | Annual Temperature Range | Sealed Road Length | Elevation | Isothermality | Surface Area of Bodies of Water | Disease Prevalence |
|---|---|---|---|---|---|---|---|---|---|
| CANX | | | X | X | | | | X | |
| FGGY | X | | | | | | | | X |
| MGAT4A | | X | | X | XX | XX | X | | |
| ARPC2 | | | | | | XX | | | |
| SCN9A | | | | X | XX | XX | | | |
| TBXAS1 | XX | | | | | | | X | X |
| ARMC3 | X | | | | X | | | | |

**Supplementary Figure 1**: Population assignments computed by fastSTRUCTURE for samples collected prior to (a,c) and post (b,d) DFTD arrival. Each vertical bar represents a single individual sampled at one of the sampling locations which are abbreviated along the x-axis. (a) Pre-DFTD plot depicting K=9 pre-DFTD arrival and (b) K=8 post-DFTD arrival determined by maximum marginal likelihood. Plots depicting K=4 (c) pre-DFTD and (d) post-DFTD. Each color represents a distinct genetic cluster.

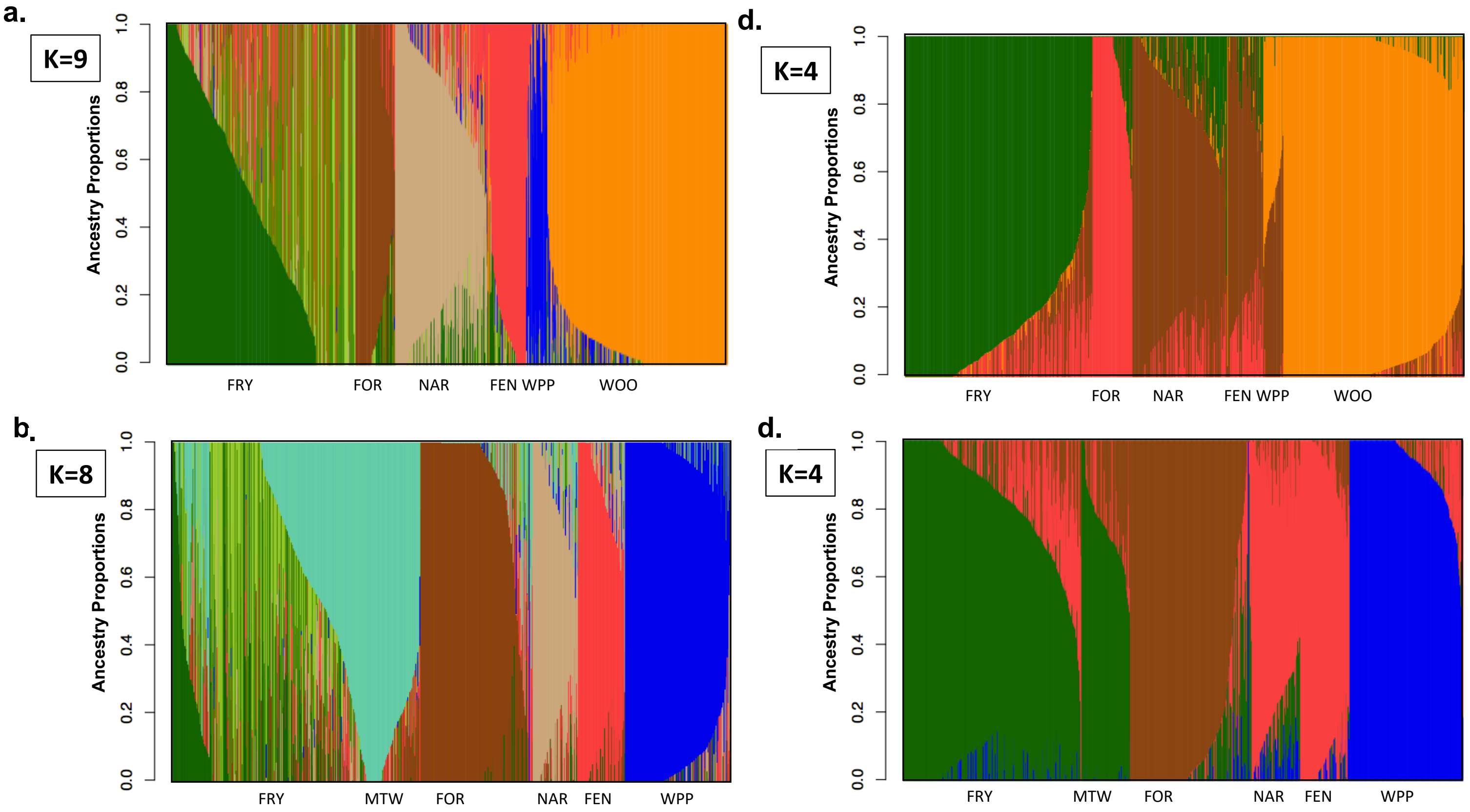

**Supplementary Figure 2**: Venn diagram showing the overlap of MINOTUAR candidate genes for the eight abiotic environmental variables in the pre-disease (left) and post-disease (right) candidate gene sets.

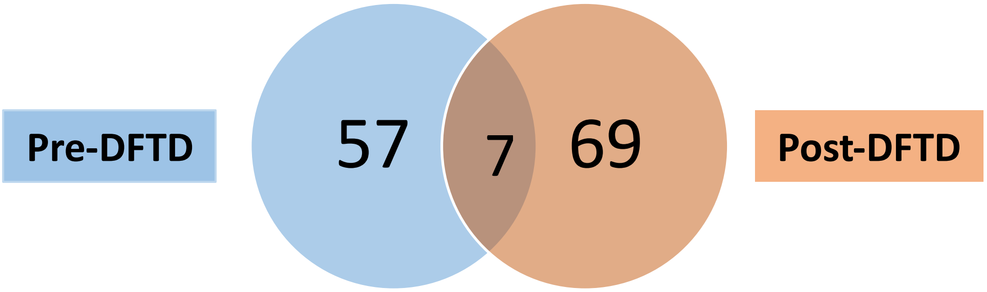

**Supplementary Figure 3**: (a-g) Mahalanobis distances from MINOTAUR for each SNP pre-DFTD arrival (left plot) and post-DFTD arrival (right plot) for seven of the abiotic environmental variables. SNPs are ordered by position along the chromosomes. The top 1% of loci with the largest Mahalanobis distance values pre-DFTD are indicated in red. (h-n) More pre-DFTD candidate loci were not identified as candidates post-DFTD than would be expected by random chance (Fisher's exact tests).

a.

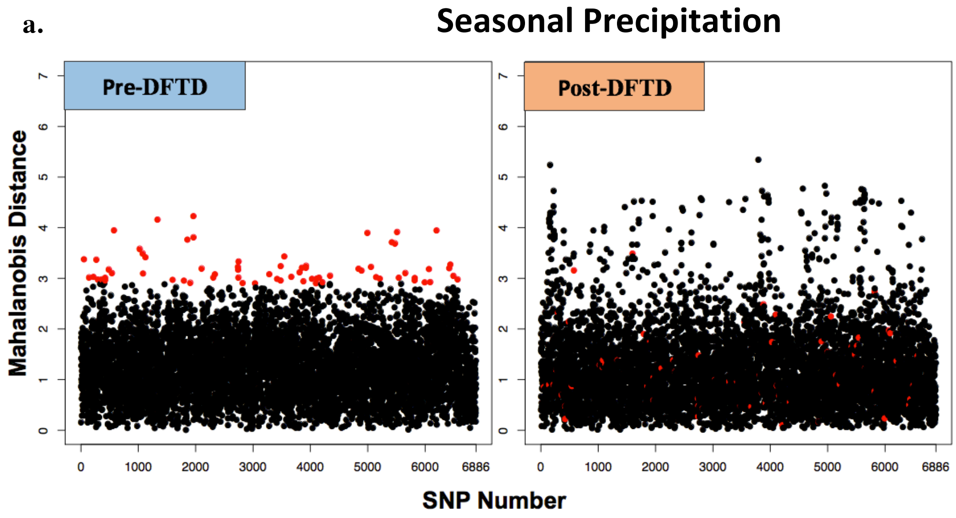


h.

| | Pre-DFTD | Post-DFTD |
|---|---|---|
| Candidate SNPs | 69 | 0 |
| Non-Candidate SNPs | 6817 | 6886 |

Odds Ratio = N/A
P-value < 2.2e-16

b.

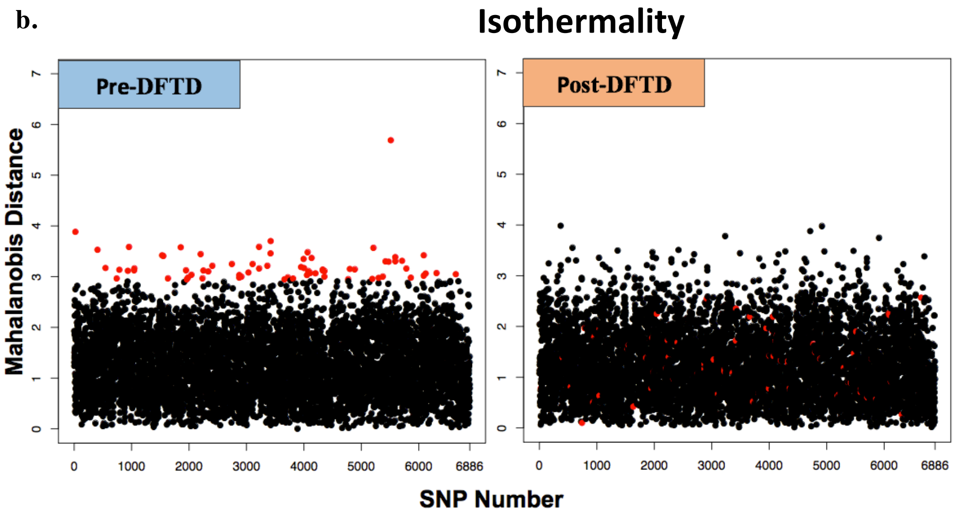


i.

| | Pre-DFTD | Post-DFTD |
|---|---|---|
| Candidate SNPs | 69 | 0 |
| Non-Candidate SNPs | 6817 | 6886 |

Odds Ratio = N/A
P-value < 2.2e-16

c. **Annual Temperature Range**

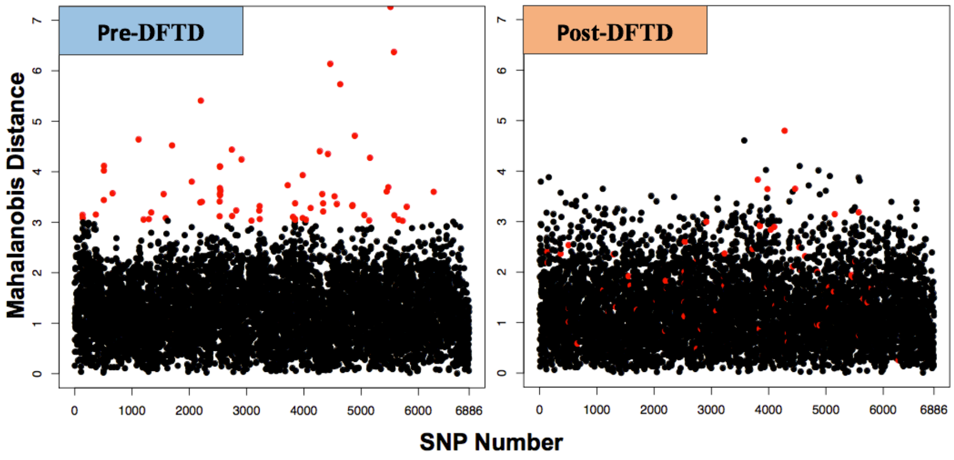


j.

| | Pre-DFTD | Post-DFTD |
|---|---|---|
| **Candidate SNPs** | 69 | 3 |
| **Non-Candidate SNPs** | 6817 | 6883 |

**Odds Ratio = 11.60**
**P-value < 5.04e-15**

d. **Vegetation Index**

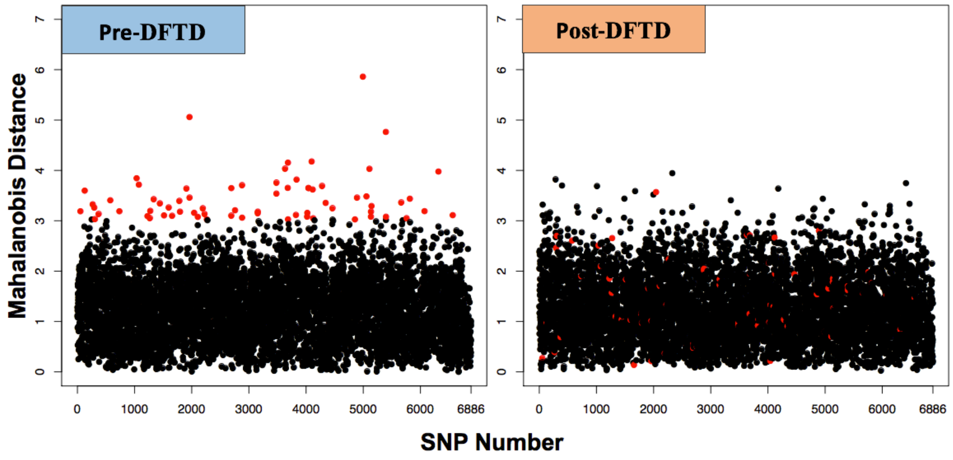


k.

| | Pre-DFTD | Post-DFTD |
|---|---|---|
| **Candidate SNPs** | 69 | 1 |
| **Non-Candidate SNPs** | 6817 | 6887 |

**Odds Ratio = 68.67**
**P-value < 2.2e-16**

e. **Length of Sealed Roads**

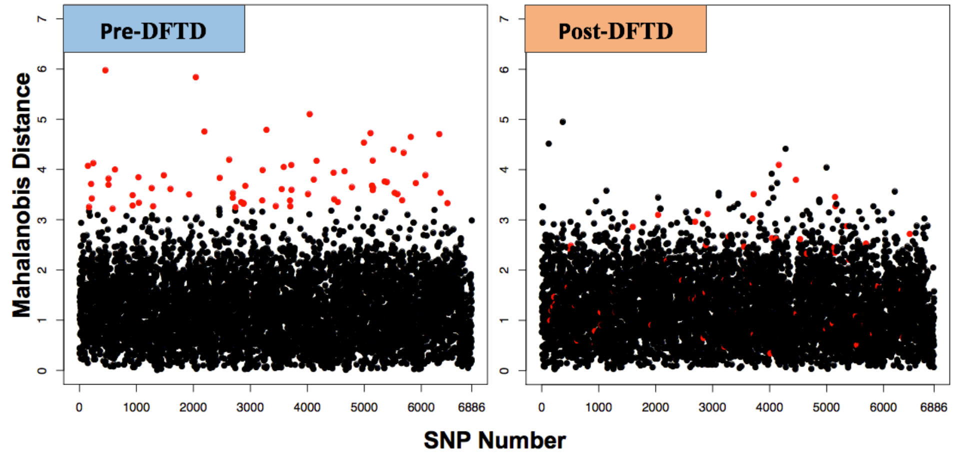


l.

| | Pre-DFTD | Post-DFTD |
|---|---|---|
| **Candidate SNPs** | 68 | 9 |
| **Non-Candidate SNPs** | 6817 | 6877 |

**Odds Ratio = 7.73**
**P-value < 2.2e-16**

f.

### Elevation

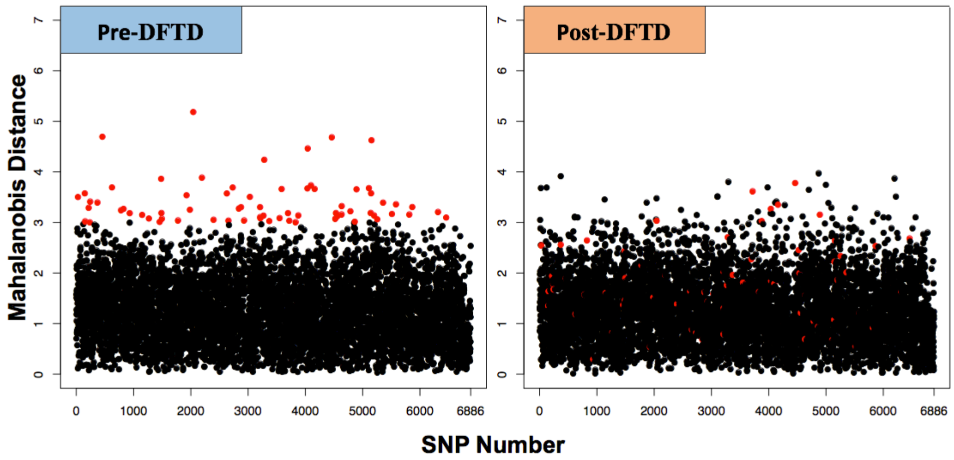


m.

| | Pre-DFTD | Post-DFTD |
|---|---|---|
| Candidate SNPs | 69 | 7 |
| Non-Candidate SNPs | 6817 | 6879 |

**Odds Ratio = 9.95**
**P-value < 2.7e-14**

g.

### Surface Area of Bodies of Water

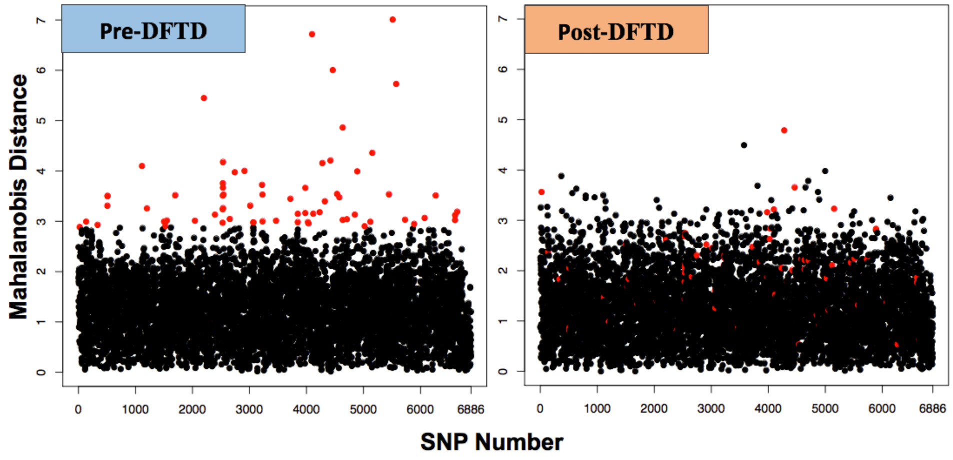


n.

| | Pre-DFTD | Post-DFTD |
|---|---|---|
| Candidate SNPs | 69 | 6 |
| Non-Candidate SNPs | 6810 | 6880 |

**Odds Ratio = 11.60**
**P-value < 5.04e-15**
